# Biomarker-Targeted O^6^-Guanine Alkylation Potentiates ATR Inhibitor Response in De Novo and Relapsed/Refractory AML

**DOI:** 10.64898/2026.09.19.751270

**Authors:** Prateek Bhardwaj, Amro Baassiri, Ranjini Sundaram, Sam Friedman, James Elia, Jennifer VanOudenhove, Collin Heer, Ahmad Kiwan, Martin Matthews, Susan Gueble, Stephanie Halene, Ranjit Bindra

## Abstract

Acute myeloid leukemia (AML) remains limited by high relapse rates and a scarcity of biomarker-directed therapies, underscoring the need to identify actionable vulnerabilities and mechanisms of therapeutic resistance. O^6^-methylguanine-DNA methyltransferase (MGMT), a DNA repair enzyme that directly reverses mutagenic O^6^-alkylguanine lesions, is epigenetically silenced in multiple cancers. Although MGMT silencing is predictive of temozolomide (TMZ) response in glioblastoma with significant survival advantage, prior clinical trials of TMZ in AML have shown only modest responses. Therefore, the prevalence and therapeutic relevance of MGMT silencing, as well as genetic factors that modify its therapeutic response in AML, remain to be comprehensively investigated. Here, we first profiled MGMT status across a diverse cohort of 23 de novo and 16 relapsed/refractory (R/R) primary AMLs using an integrated analysis of MGMT mRNA expression, promoter methylation, and protein expression. We found that approximately 25-30% of AMLs harbor MGMT silencing, in contrast to consistent MGMT expression in healthy CD34^+^ hematopoietic stem and progenitor cells. We further identified frequent loss of mismatch repair (MMR) proteins in these AML cohorts, a known resistance mechanism against TMZ in glioblastoma. Using CRISPR knockout screening and genetically engineered AML models, we found that MMR loss also drives both upfront and acquired TMZ resistance in MGMT-silenced AML. These findings identify MMR deficiency as an important and previously underappreciated genetic contributor to the modest responses observed in prior trials of TMZ in AML. To therapeutically exploit MGMT silencing while bypassing genetic determinants of resistance, including MMR deficiency, we evaluated clinically and preclinically characterized MGMT-dependent DNA-alkylating agents and identified a fluoroethylating analog of TMZ, N^3^-(2-fluoroethyl) imidazotetrazine (KL50), with pronounced and selective activity in MGMT-silenced AML. KL50 retained antileukemic activity irrespective of MMR status, significantly prolonging survival in humanized MISTRG6 mice harboring primary AML patient-derived xenografts. Mechanistically, KL50 induced DNA damage through the time-dependent formation of interstrand DNA crosslinks, bypassing MMR dependence and triggering a replication stress response dominated by ATR signaling. Pharmacologic ATR inhibition synergized with KL50, producing marked antileukemic activity and significantly extending survival across AML models without compromising hematologic safety. Together, these findings establish MGMT silencing as a prevalent and therapeutically actionable biomarker in AML, define MMR status as a key determinant of TMZ response in AML but not KL50 sensitivity, and provide a translational rationale for combining low-dose O^6^-fluoroethylating imidazotetrazines with ATR inhibitors to target both de novo and R/R AML across diverse genetic backgrounds.

## Introduction

Acute Myeloid Leukemia (AML) is a heterogeneous hematologic malignancy defined by clonal expansion of poorly differentiated myeloid progenitors and impaired hematopoiesis ^1^. Despite recent advances in treatment with integration of targeted agents into chemotherapy, low-intensity treatments as well as stem cell transplantation, therapeutic failure and relapse remain frequent ^2^. Defining additional molecular vulnerabilities is therefore critical to expand therapeutic strategies for patients with high-risk and relapsed/refractory (R/R) AML.

O^6^-methylguanine-DNA methyltransferase (MGMT) silencing through promoter methylation represents a recurrent DNA damage response vulnerability across cancer types, including AML ^3^. MGMT encodes a 22 kDa repair protein that removes alkyl adducts from the mutagenic O^6^-guanine position by transferring the lesion to cysteine-145 within its active site, resulting in irreversible inactivation and proteasomal degradation ^4^. Because each MGMT molecule repairs a single lesion, sustained MGMT expression is required to preserve repair capacity. Loss or reduction of MGMT permits persistence of O^6^-methylguanine lesions, thymine mispairing during replication, and accumulation of G>A and C>T transition mutations, thereby promoting genomic instability and clonal evolution ^5–7^. MGMT promoter methylation is an established predictive biomarker for temozolomide (TMZ) response in glioblastoma ^8^. AML and glioma share alterations in epigenetic regulatory pathways, including IDH1/2-associated metabolic and DNA methylation changes, providing a biological rationale for investigating the therapeutic relevance of MGMT silencing in AML ^9–11^, which is insufficiently characterized despite a reported prevalence of approximately 25-30% ^5^. Two clinical studies have reported response rates of approximately 40-60% among MGMT-deficient AML patients treated with TMZ, although with limited durability ^12,13^. Intact mismatch repair (MMR), which converts mispaired O^6^-methylguanine lesions into replication-associated DNA damage through futile repair cycling, is required for TMZ toxicity in MGMT-deficient cells ^14^. Intrinsic or acquired MMR loss represents a major mechanism of TMZ resistance in solid tumors ^14–20^. This resistance axis may be particularly relevant in AML, where structural or functional MMR defects have been reported to increase from approximately 21% at diagnosis to nearly 48% in heavily treated R/R disease ^21^. Additional alkylating agents belonging to the triazene, nitrosourea, and methane sulfonate classes can generate adducts at O^6^-guanine. Agents that retain strong MGMT-selectivity without susceptibility to MMR-mediated resistance could provide a therapeutic opportunity, in particular for R/R MGMT-deficient AML ^22,23,24^.

We validate MGMT loss in approximately 25-30% of both de novo and R/R AML and that MMR deficiency indeed functions as an important determinant of resistance to TMZ. Whole-exome sequencing of single-cell-derived clonal tumors further demonstrates that chronic TMZ exposure rapidly selects for MMR-deficient, hypermutated subpopulations. We previously showed that KL-50, a fluoroethyl imidazotetrazine analog of TMZ, generates O^6^-fluoroethylguanine lesions that mature into interstrand DNA crosslinks ^22^. Here, we demonstrate that, similar to solid tumors, KL-50-induced crosslinks exert cytotoxicity in AML independently of the MMR-mediated futile cycling required for TMZ activity. KL-50-induced crosslinks generate replication stress, creating a dependency on ATR signaling. Although ATR inhibitors are already in clinical development, their therapeutic application can be constrained by limited biomarker selection and hematologic toxicity ^25,26^. We show that sequential O^6^-fluoroethylation followed by ATR inhibition provides a selective and tolerable therapeutic strategy for MGMT-deficient AML.

## Results

### MGMT and MMR silencing is highly prevalent across distinct subsets of de novo and relapsed/refractory (R/R) AML

Glioma and AML share prominent epigenetic vulnerabilities, with recurrent IDH1 mutations in glioma and alterations involving IDH1/2, TET2, and DNMT3A in AML ^27,28^. MGMT silencing in glioma is most commonly associated with promoter hypermethylation ^29^. IDH mutations provide a broader epigenetic context for this phenotype through production of 2-HG and inhibition of α-KG-dependent demethylation, resulting in extensive DNA hypermethylation and establishment of Glioma CpG Island Methylator Phenotype (G-CIMP) ^9,11^. MGMT expression can additionally be influenced by enhancer and higher-order chromatin regulation, histone modifications, transcriptional regulation, microRNAs, and post-transcriptional/post-translational mechanisms ^30^. Analysis of TCGA transcriptomic data confirmed that AML, together with several glioma subtypes, is among the lowest MGMT-expressing tumor types (Supplementary Figure S1A) ^5^. Meta-analysis of the BloodSpot dataset further suggested low expression in leukemic cells compared with normal hematopoietic stem and progenitor cells (Supplementary Figure S1B). As MGMT mRNA expression has historically shown imperfect correlation with protein abundance, we measured MGMT protein levels in 39 AML samples, comprised of 23 de novo and 16 R/R AMLs, and in mobilized CD34^+^ progenitors from three healthy donors. MGMT protein levels were low to absent in 26% (6/23) of de novo and 43.7% (7/16) of R/R AMLs (Figure 1A-C, Supplementary Table T1). We also observed loss or barely detectable levels of one or more MMR proteins, such as MSH2, MSH6, MLH1, or PMS2 in 65% (15/23) of de novo and 62% (10/16) of R/R AMLs (Figure 1A). Reduced MMR expression was also evident in the meta-analysis of the BloodSpot datasets, suggesting that MMR loss may be a frequent feature of AML (Supplementary Figure S1C-E). At least four de novo (4/23) and six R/R (6/16) samples exhibited concurrent low MGMT and MMR protein levels, demonstrating frequent co-occurrence of these two phenotypes. Consistent with the known requirement for functional MMR in TMZ cytotoxicity, a representative MGMT-low/MMR-low sample (Y1063) was insensitive to TMZ, similar to the MGMT-proficient AML sample Y2315 and healthy CD34^+^ cells, whereas the MGMT-low/MMR-proficient sample Y4008 showed TMZ-sensitivity (Figure 1D). This mirrors the intrinsic TMZ resistance associated with MMR deficiency in glioblastoma and suggests that MMR status could have been an underappreciated confounder in prior MGMT-focused clinical studies of TMZ in AML ^12,13^.

**Figure 1:**
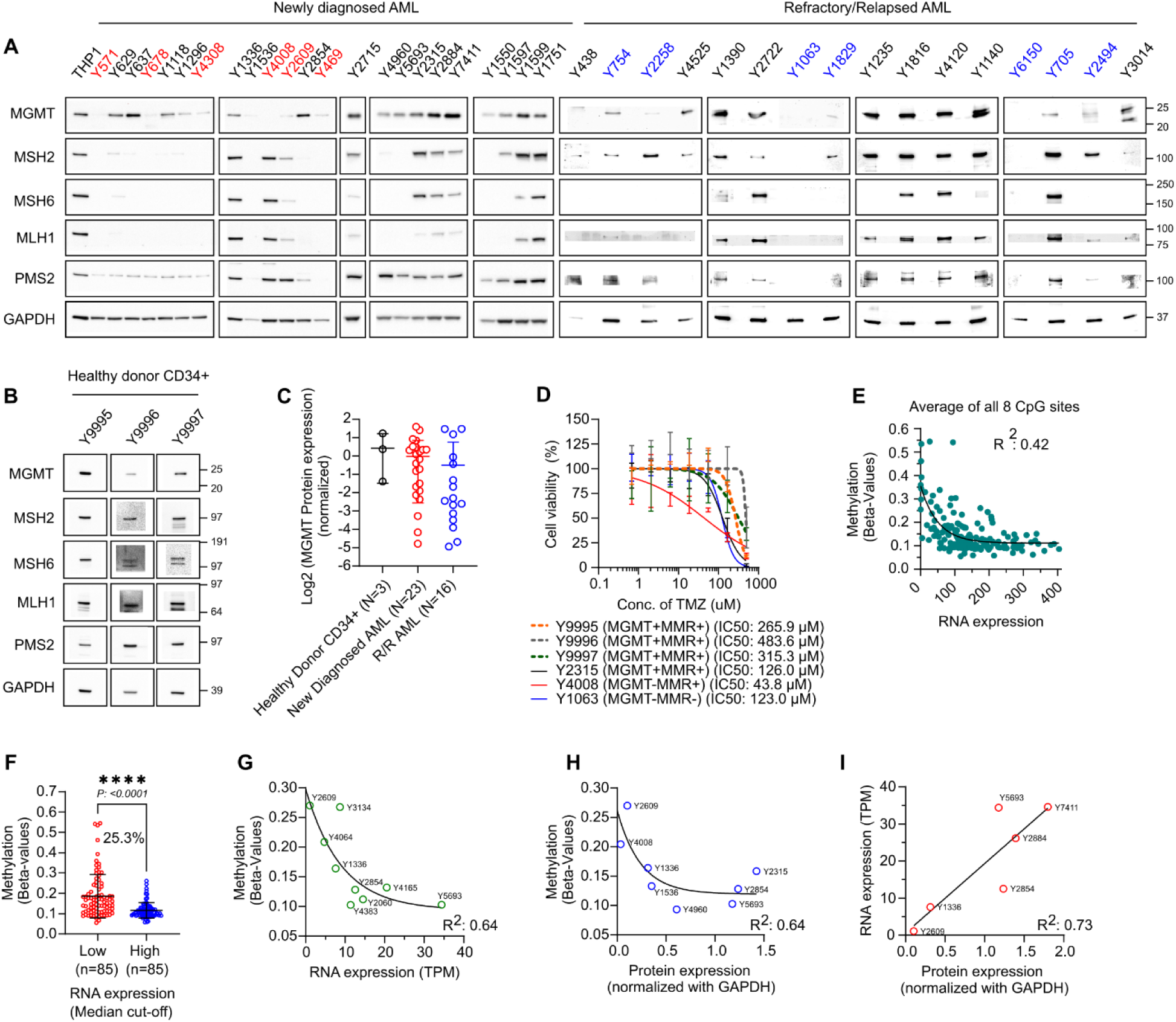
Western blot analysis of MGMT and core MMR proteins (MSH2, MSH6, MLH1, and PMS2) in **A)** de-identified newly diagnosed and R/R primary AML samples (red and blue highlights denote MGMT-low samples) and **B)** CD34+ cells from healthy donors from the Yale Hematology Biobank. **C)** Normalized MGMT protein expression quantified via Western blot showed in A and B in healthy donor CD34+ cells, compared with bone marrow samples from newly diagnosed and R/R AML patients. MGMT expression was normalized to GAPDH (loading control) and the THP1 cell line (chemiluminescence exposure control) across all blots. **D)** 72 hour *in vitro* viability assays for TMZ in de-identified MGMT^-^MMR^-^ R/R AML (Y1063), MGMT^-^MMR^+^ AML (Y4008), MGMT^+^MMR^+^ AML (Y2315), and healthy CD34^+^ cells. **E)** Correlation analysis between MGMT mRNA expression and mean DNA methylation across all 8 significantly negatively correlated promoter CpG sites in the TCGA AML cohort (n=170). **F)** Comparison of mean methylation (beta-values of eight selected CpG probes) between MGMT-low and MGMT-high AML patients from TCGA dataset, stratified by median expression. **G-H)** Correlation of mean methylation with **G)** mRNA expression (n=9) and **H)** protein expression (n=8) in matched patient samples from the Yale Heme-Biobank. Goodness of fit was determined using a one-phase exponential decay non-linear regression model. **I)** Linear regression analysis correlating mRNA and protein expression in matched AML samples (n=6).

MGMT promoter methylation serves as biomarker of TMZ response in glioma; we next investigated whether promoter methylation similarly explained MGMT silencing in AML. We identified eight CpG sites within the MGMT promoter region (-552 to +289 bp relative to the transcription start site) in the TCGA AML cohort ^31^ that showed the strongest inverse correlations with MGMT RNA expression across 170 patients (Supplementary Figure S2A-B, Figure 1E). To estimate the frequency of epigenetic MGMT silencing, we stratified the TCGA cohort into MGMT-low and MGMT-high groups using median mRNA expression as the cutoff. The MGMT-low group exhibited significantly greater methylation across these eight CpG sites, with approximately 25% of patients showing methylation levels greater than one standard deviation above the mean of the MGMT-high group (Figure 1F). We next measured methylation at these CpG sites and MGMT mRNA expression in our primary AML samples and examined their concordance with MGMT protein levels. Methylation across the eight CpG sites inversely correlated with both MGMT mRNA and protein abundance, and MGMT mRNA and protein levels showed a strong positive correlation (Figure 1G-I). Collectively, these independent analyses across the TCGA and Yale cohorts establish concordance among MGMT promoter methylation, transcriptional suppression, and reduced protein expression, supporting epigenetic silencing as a major mechanism underlying MGMT deficiency in a molecularly defined subset of AML.

### MGMT-silenced AML models exhibit selective vulnerability to O^6^-fluoroethylating imidazotetrazine

TMZ-induced O^6^-methylguanine cytotoxicity requires functional MMR. We curated a structurally diverse library of mono- and bifunctional alkylating agents that target DNA nucleophilic sites with varying propensities, including N^7^-guanine, N^3^-adenine, and O^6^-guanine. Unlike N^7^-guanine and N^3^-adenine lesions, which are predominantly repaired through base excision repair, O6-guanine lesions are directly repaired by MGMT, providing a basis for MGMT-dependent selectivity ^24,32–34^. The library included imidazotetrazines (TMZ, MTZ, and 2-fluoroethyl analog of TMZ), nitrosoureas (lomustine and carmustine), and methyl methanesulfonate (MMS), along with clinically used N7-guanine crosslinkers (busulfan, cyclophosphamide, chlorambucil, and bendamustine) as controls. To quantify MGMT-dependent selectivity, we calculated a therapeutic index (T.I.), defined as the ratio of IC50 in MGMT-proficient to IC50 in MGMT-deficient isogenic cells. Across MGMT-isogenic pairs of U937 and MOLM13 cells, imidazotetrazines showed the highest T.I. values, with KL50 emerging as the leading hit with a T.I. comparable to that of TMZ (Figure 2A, Supplementary Figure S3A). We confirmed favorable TMZ and KL-50 T.I.s across an extended panel of isogenic myeloid leukemia cell lines (Figure 2B, Supplementary Figure S3B). To orthogonally validate these findings, we analyzed KL50 efficacy across 22 AML cell lines from a comprehensive 900-cell-line PRISM (Profiling Relative Inhibition Simultaneously in Mixtures) screen conducted with the Broad Institute ^35^. MGMT expression emerged as the strongest predictor of KL50 efficacy in AML. In contrast, the association between MGMT expression and TMZ response failed multiple-testing correction (q = 0.997), consistent with additional determinants of TMZ sensitivity beyond MGMT alone ^33^ (Figure 2C-D). To define the relationship between MGMT abundance and KL50 sensitivity, we generated isogenic clones expressing graded levels of MGMT protein in MGMT-deficient MOLM13 cells (Supplementary Figure S3C). KL50 response showed a sigmoidal relationship with MGMT expression similar to TMZ, characterized by an initial lag phase followed by a steep increase in resistance beyond a threshold MGMT level and a plateau at higher MGMT expression, potentially reflecting an increasing contribution of MGMT-independent lesions. (Figure 2E, Supplementary Figure S3D-F).

**Figure 2.**
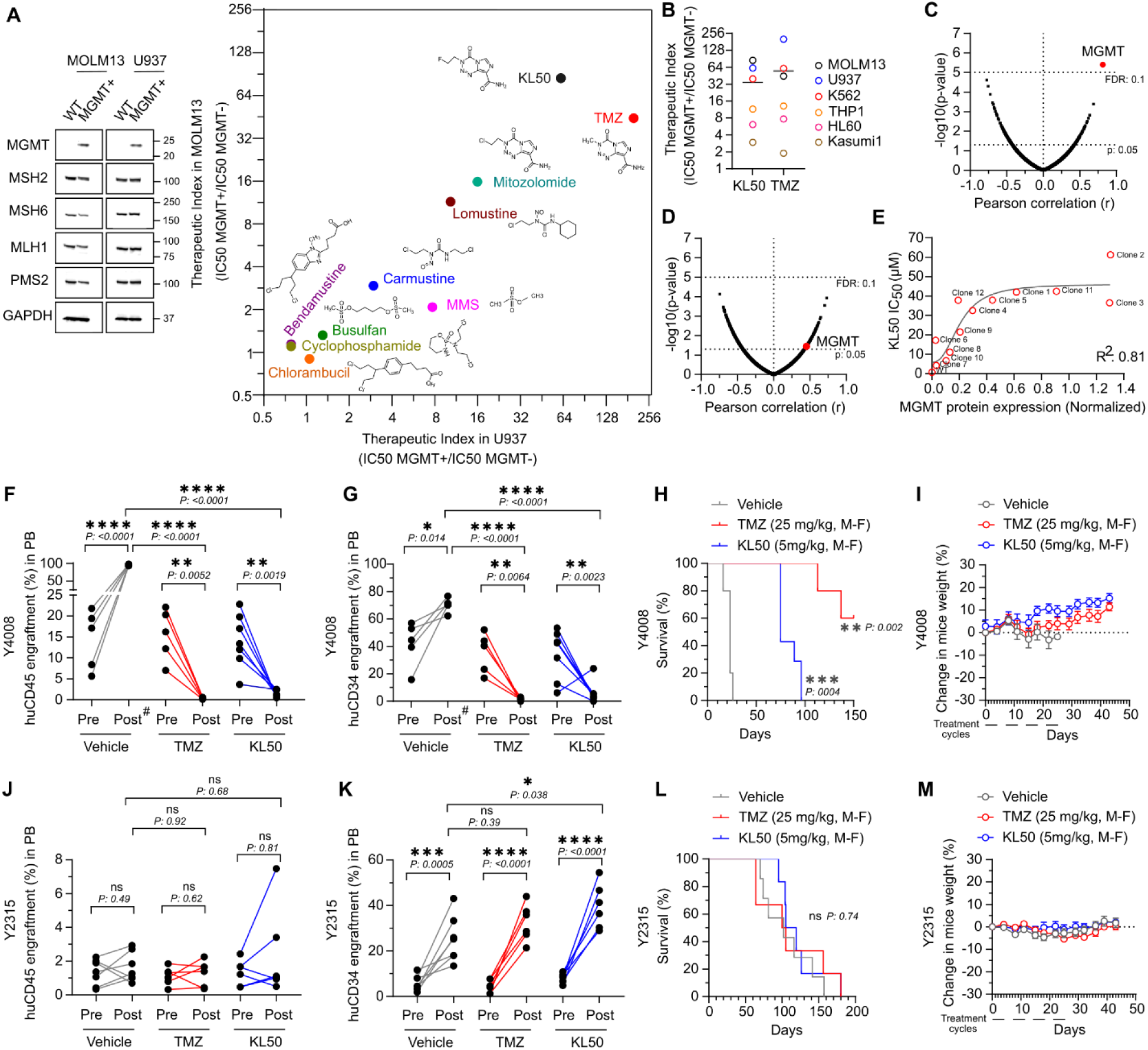
**A)** Western blot confirming MGMT ORF expression in wild-type (MGMT-null) MOLM13 and U937 cells alongside Therapeutic Index (T.I.) of diverse alkylating agents in MGMT-isogenic pairs of MOLM13 and U937 cells. T.I. is defined as the ratio of IC50 in MGMT^+^ cells to the IC50 in MGMT^-^ counterparts. **B)** T.I. of KL50 and TMZ across a panel of MGMT-isogenic AML cell lines. **C-D)** Volcano plots correlating gene expression (n = 15,971 genes) with sensitivity to **C)** KL50 and **D)** TMZ across 22 AML cell lines from the PRISM screen. **E)** Correlation between MGMT protein levels and sensitivity to KL50 in isogenic MOLM13 clones. Goodness of fit was determined using a sigmoidal 4PL non-linear regression model. **F-M)** *In vivo* efficacy of KL50 and TMZ in MISTRG6 mice engrafted with **F-I)** MGMT^-^ (Y4008) and **J-M)** MGMT^+^ (Y2315) primary AML samples. “Pre” and “Post” in Panel F,G,J,K denote peripheral blood (PB) engraftment levels at Day 0 and Day 42 (two weeks post-completion of 4 treatment cycles), respectively. **H,L)** Kaplan-Meier survival curves and **I,M)** body weights of MISTRG6 mice engrafted with Y4008 and Y2315 under TMZ and KL50 treatment. Treatment cycle: 5 days a week (M-F, Monday to Friday) for 4 weeks. (#) in panel F and G denotes that the CD45^+^ and CD34^+^ engraftment analysis represents the 2-week timepoint from study initiation, as the vehicle group began dropping out due to high tumor burden as reflected in the survival curve (H).

To determine whether KL50 shows efficacy *in vivo*, we tested KL50 in cell line derived xenografts (CDXs) of MGMT-deficient, MMR-proficient U937 cells. KL50 demonstrated efficacy comparable to TMZ (Supplementary Figure S4A). We next evaluated KL50 in primary AML xenografts in cytokine-humanized immunodeficient mice. Adult MISTRG6 were engrafted with primary MGMT-high (Y2315) or MGMT-low (Y4008) AML. Mice were assigned to treatments cohorts in balanced fashion based on leukemia burden in peripheral blood (by huCD45^+^ and huCD34^+^ percentages) and hematologic parameters (platelet counts) (Supplementary Figure S4B-F). KL50 and TMZ demonstrated pronounced MGMT-selective activity, resulting in significant reductions in huCD45^+^ and huCD34^+^ leukemic populations and significantly extending survival in mice harboring the MGMT-low Y4008 PDX, but not the MGMT-high Y2315 PDX (Figure 2F-H, 2J-L). Both agents were well tolerated without significant weight loss with improvement in platelet counts and hemoglobin in Y4008 engrafted mice and no changes in Y2315 engrafted mice (Figure 2I,M, and Supplementary Figure S4G-J). Collectively, these *in vivo* findings in humanized primary AML xenografts validate the antileukemic efficacy and MGMT selectivity of KL50.

### O^6^-fluoroethylating imidazotetrazine KL50 retains activity independent of mismatch repair status in MGMT-deficient AML

Fluoroethylating imidazotetrazines such as KL50 lack MMR dependence in solid tumor models ^22^. To systematically identify possible genetic modifiers of KL50 response in AML, we performed a focused pooled CRISPR-knockout screen targeting 353 core DNA damage response (DDR) genes, including all major MMR components, in MOLM13 cells. Following two pulses of KL50 at an IC30 dose, gRNAs targeting Fanconi anemia (FA) and homologous recombination (HR) genes were significantly negatively enriched, suggesting that MGMT-deficient AML cells depend on these pathways to tolerate or repair KL50-induced DNA damage. TMZ similarly produced synthetic-lethal interactions with several of these DDR pathways (Figure 3A and Supplementary Figure S5A,C). When selection pressure was increased to IC87, loss of the major MMR genes MSH2, MSH6, MLH1, and PMS2 resulted in positive enrichment under TMZ but not KL50 treatment (Supplementary Figure S5B,D). In contrast, loss of TP53 and CHEK2 emerged among the strongest resistance-associated hits for KL50. To orthogonally validate the MMR independence of KL50, we depleted MSH6 using shRNA, as disruption of a single component of the heterodimeric MMR complexes: MutSα (MSH2-MSH6) or MutLα (MLH1-PMS2), is sufficient to compromise pathway function ^36^. MSH6 loss in three isogenic models had no significant effect on KL50 cytotoxicity (Figure 3B-D). In contrast, MSH6 loss markedly increased TMZ IC50 and reduced its *in vitro* efficacy (Supplementary Figure S5E-J), and even a moderate 45–50% reduction in MSH6 protein was associated with an approximately 73-fold increase in TMZ IC50 (Supplementary Figure S5E-K). MMR independence was confirmed for KL50, whereas TMZ lost efficacy in MSH6-deficient CDXs (Figure 3E-H and Supplementary Figure S5L-M). Finally, KL50 retained *in vitro* activity in an MGMT-deficient/MMR-deficient primary AML (Y1063), comparable to that observed in the MGMT-deficient/MMR-proficient Y4008 AML (Figure 3I).

**Figure 3:**
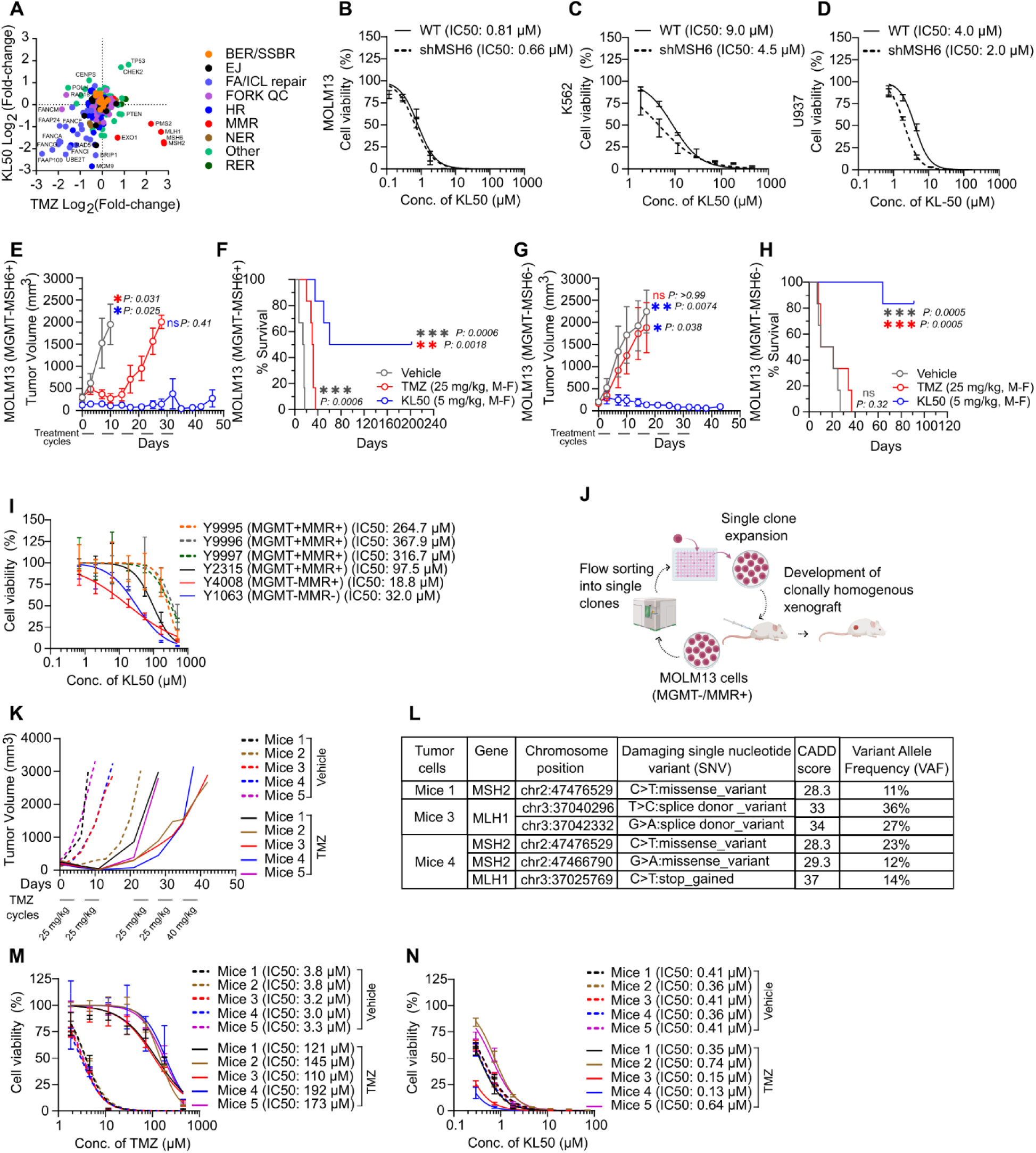
**A)** Comparative plot of gRNA enrichment and dropout from the DDR-focused CRISPR-knockout screen in parental MGMT^-^ MOLM13 cells treated with 1.84 µM KL50 and 17 µM TMZ. **B-D)** Short-term (6-day) *in vitro* viability assays with KL50 in MSH6-isogenic AML models following knockdown in **B)** MOLM13 (shMSH6 clone 5), **C)** K562 (shMSH6 clone 4), and **D)** U937 (shMSH6 clone 10). **E-H)** *In vivo* tumor growth inhibition and Kaplan-Meier survival analysis of Athymic Nude-Foxn1nu mice bearing **E-F)** MGMT^-^MMR^+^ and **G-H)** MGMT^-^MMR^-^ MOLM13 subcutaneous xenografts. **I)** 72 hour *in vitro* viability assays with KL50 on de-identified MGMT^-^MMR^-^ R/R AML (Y1063), MGMT^-^MMR^+^ AML (Y4008), MGMT^+^MMR^+^ AML (Y2315), and healthy CD34^+^ cells (Y9995, Y9996, Y9997). **J)** Schematic showing development of clonally homogenous MOLM13 xenograft in Nude mice. **K)** *In vivo* tumor growth kinetics in a monoclonal MOLM13 xenograft nude mice model under escalating TMZ dosage. **L)** Table showing genomic coordinates, type of variant, allele frequency, and Combined Annotation Dependent Depletion (CADD) score of the damaging variant alleles detected in the MMR genes of MOLM13 tumor cells harvested from mice 1, 3, and 4 of the TMZ-treated cohort. **M-N)** Short-term *in vitro* viability assays on harvested tumor cells from Vehicle and TMZ-treated cohorts comparing sensitivity to **M)** TMZ and **N)** KL50.

We next asked whether KL50 retains activity in AML tumors that acquire resistance to TMZ. In glioma and other solid tumors, prolonged TMZ exposure can induce a hypermutator phenotype with clonal selection of mutations in MMR genes ^15,20,37–39^. MGMT-deficient/MMR-proficient parental MOLM13 xenografts initially responded to TMZ but progressed despite continued treatment after only a few cycles, whereas KL50 maintained prolonged tumor control (Figure 3E). Therefore, we generated a single-cell-derived MOLM13 clonal xenograft to minimize pre-existing cellular heterogeneity (Figure 3J) that showed the same initial response to TMZ followed by rapid loss of sensitivity within 2-3 treatment cycles (Figure 3K). Tumors from vehicle- and TMZ-treated mice were harvested, dissociated into single-cell suspensions, and analyzed for MGMT and MMR protein expression. Whole-exome sequencing was performed on tumors from three representative TMZ-treated mice. TMZ-treated tumors showed reduced MLH1, MSH2, and MSH6 protein levels and acquired damaging mutations in MSH2 and MLH1 across all three analyzed tumors (Supplementary Figure S6A, and Figure 3L) with increased variant allele fractions (VAFs) relative to the overall genomic mutational background, consistent with the expansion of MMR-deficient clones under TMZ selection pressure (Supplementary Figure S6B). As expected, leukemic cells isolated from TMZ-treated mice were resistant to TMZ *in vitro*, whereas cells recovered from vehicle-treated tumors remained sensitive (Figure 3M). In contrast, KL50 retained pronounced activity against cells from both vehicle- and TMZ-treated tumors (Figure 3N). These results were further validated *in vitro* via prolonged treatments of MOLM13 and U937 cells with emergence of resistance to TMZ (Supplementary Figure S6C), reduced MMR protein expression particularly of MSH2 and MSH6 (Supplementary Figure S6D), concurrent acquired damaging mutations, clonal selection evident via elevated VAFs (Supplementary Figure S6E-F), and retained sensitivity to KL50 (Supplementary Figure S6G). Collectively, these findings demonstrate that unlike TMZ KL50 activity in AML is primarily determined by MGMT status and is retained when MMR pathway is compromised. These data nominate KL50 as valuable biomarker-driven targeted therapeutic with improved resistance profile over TMZ.

### KL50 induces an MGMT-selective replication stress dependency that can be exploited by ATR inhibition

Single-agent regimens rarely show prolonged efficacy, and combination regimens can increase cytotoxicity while mitigating adverse effects from either agent alone. We therefore sought to identify a biomarker-driven combination strategy utilizing KL50. KL50, but not TMZ, induced a significant increase in □H2AX in MGMT-deficient MOLM13 and U937 cells irrespective of MMR status (Figure 4A, Supplementary Figure S7A). Cell-cycle-resolved analysis further showed that KL50-induced □H2AX predominantly accumulated during S and G2 phases, whereas TMZ-associated damage was enriched primarily in G1 and S phases (Supplementary Figure S7B). KL50-induced O^6^-fluoroethylguanine lesions are known to mature into interstrand DNA crosslinks (ICLs), whose processing requires coordinated activity of the Fanconi anemia (FA) and homologous recombination (HR) pathways, particularly during DNA replication ^22,35^. The accumulation of KL50-induced damage in S/G2 aligned with the strong FA and HR synthetic-lethal signatures identified in our CRISPR-knockout screen (Figure 3A). Since ICL processing and stalled replication forks generate single-stranded DNA intermediates, we measured phosphorylation of RPA32 at Ser33 as a marker of replication stress. KL50 also induced robust pRPA32 in MGMT-deficient cells irrespective of MMR status (Figure 4B). Persistent replication stress and single-stranded DNA primarily activate ATR signaling, whereas DNA double-strand break-associated signaling can engage ATM. Consistent with activation of both DDR arms during processing of KL50-induced lesions, KL50 produced strong MGMT-dependent but MMR-independent phosphorylation of CHK1 (Ser345) and CHK2 (Thr68) (Figure 4C). Resulting apoptosis was verified by cleavage of PARP1 and activation of caspases 3/7 (Figure 4D).

**Figure 4:**
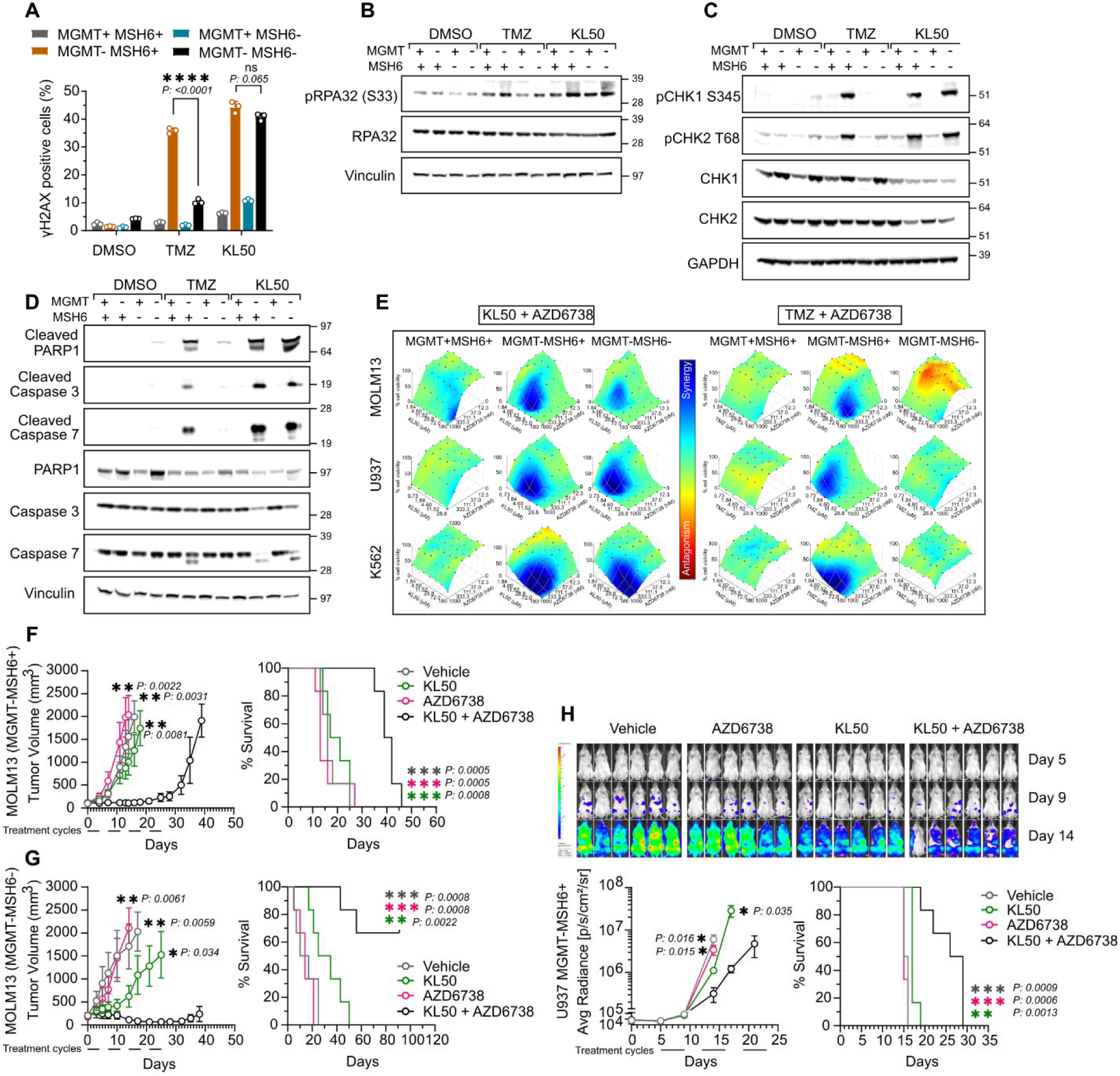
**A-D)** Comparative DNA damage and response signaling analysis post TMZ and KL50 treatment of MGMT and MMR-isogenic MOLM13 cells. Flow cytometry-based □H2AX quantitation illustrates **A)** DNA damage, and representative western blots illustrate **B)** replication stress (p-RPA32 S33), **C)** signal transduction through p-CHK1 (S345) and p-CHK2 (T68), and **D)** apoptotic induction (cleaved Caspase-3 and cleaved PARP1). **E)** Synergy evaluation of the AZD6738 (ATRi) in combination with KL50 in comparison to TMZ across a panel of AML cell lines isogenic for MGMT and MMR. **F-G)** *In vivo* synergy of KL50 and ATRi assessed via tumor growth kinetics and Kaplan-Meier survival analysis in athymic nude mice bearing subcutaneous MGMT^-^ MOLM13 xenografts isogenic for MMR. **H)** Representative bioluminescence imaging (BLI), corresponding quantitative flux analysis, and Kaplan-Meier survival curves demonstrating KL50 and ATRi synergy in an intravenously engrafted MGMT^-^ U937 AML model in NSG mice. Treatment cycle: Twice-weekly alternating doses (KL50, 5 mg/kg: Mon/Thu; ATRi, 50 mg/kg: Tue/Fri) for 3-4 weeks.

We next asked whether these KL50-induced DDR dependencies could be therapeutically exploited. We evaluated KL50 in combination with inhibitors targeting ATR, ATM, PARP, and CHK1, as well as agents representing major therapeutic backbones in AML, including BCL-2 inhibitors and hypomethylating agents. Among these combinations, ATR inhibition produced the most pronounced synergy with KL50, and importantly, this interaction remained strongly MGMT-dependent but MMR-independent (Supplementary Figure S7C-D and Figure 4E). These findings suggested that despite engagement of multiple DDR pathways, KL50-treated MGMT-deficient cells develop a particularly strong dependency on ATR-mediated replication stress signaling. The ATR inhibitor ceralasertib (AZD6738) has been under clinical evaluation, including studies in myeloid malignancies (NCT03770429); however, therapeutic development of ATR inhibition can be limited by hematologic toxicity ^25,26^. We therefore reasoned that KL50-induced damage in MGMT-deficient AML could provide a strategy to enhance sensitivity to lower or intermittent dosing of ATR inhibitors. Additionally, O^6^-fluoroethylguanine lesions generated by KL50 mature into ICLs with an *in vitro* half-life of approximately 18 hours ^22,35^, providing a mechanistic rationale for temporally separating lesion formation from inhibition of the subsequent ATR-dependent stress response. Indeed, exposure to KL50 followed by ATR inhibition 24 hours later showed improved synergistic cell killing across multiple AML models compared with concurrent treatment (Supplementary Figure S7E). Low-dose sequential combination *in vivo* in subcutaneous MOLM13 and intravenously engrafted U937 xenograft models, including MMR-isogenic backgrounds, produced significant tumor growth inhibition and prolonged survival irrespective of MMR status, without significant changes in body weight, supporting tolerability and biomarker-dependent efficacy of this combination (Figure 4F-H and Supplementary Figure S7F-L). Collectively, these findings identify ATR-mediated replication stress signaling as a therapeutically actionable dependency created by KL50 in MGMT-deficient AML. Temporally sequencing ATR inhibition after formation and processing of KL50-induced O6-fluoroethylguanine lesions enhances antileukemic activity while maintaining MGMT selectivity and bypassing MMR dependence, providing a mechanistically guided strategy for combining low-dose O6-fluoroethylation with ATR inhibition.

## Discussion

The therapeutic landscape of AML remains challenged by rapid clonal evolution, intrinsic treatment resistance, and frequent emergence of R/R disease ^40–42^. Although FLT3, IDH1/2, DNMT3, BCL-2, and menin inhibitors have expanded targeted treatment options for molecularly defined AML subsets, there remains a substantial need to identify additional actionable biomarkers and corresponding therapeutic strategies, particularly for R/R disease. Here, we establish epigenetic silencing of the DNA repair protein MGMT as a prevalent and therapeutically actionable vulnerability in AML, observed in approximately 25–30% of de novo and around 40% of R/R disease. We further identify MMR deficiency as an important determinant limiting the efficacy of conventional MGMT-directed methylating therapy and demonstrate that the O^6^-fluoroethylating imidazotetrazine KL50 circumvents this resistance mechanism. Finally, by defining the replication stress response generated by KL50, we identify ATR inhibition as a rational, biomarker-directed combination strategy for MGMT-silenced AML.

Historical attempts to exploit MGMT deficiency in myeloid malignancies using TMZ produced encouraging initial responses but limited overall and durable clinical benefit ^12,13^. Our findings provide a potential mechanistic explanation for this discrepancy by identifying both baseline and acquired MMR deficiency as critical modifiers of TMZ response in AML. TMZ cytotoxicity following O^6^-methylguanine formation requires functional MMR to convert persistent O^6^-methylguanine mispairs into replication-associated DNA damage through futile repair cycling ^14^. Consequently, MMR loss provides a direct route of escape from TMZ despite continued MGMT deficiency ^15^. Consistent with this mechanism, we observed frequent reduction of MMR proteins in both de novo and R/R AML, including co-occurrence with MGMT deficiency. Moreover, prolonged TMZ exposure repeatedly promoted the emergence and selection of MMR-deficient populations harboring alterations in MutSα and MutLα components across genetically distinct AML models. In some tumors, this occurred alongside independent clones exhibiting MGMT re-expression associated with reduced promoter methylation. Thus, TMZ treatment can select multiple parallel resistance routes within an initially sensitive MGMT-deficient population, providing a plausible explanation for the heterogeneous and short-lived responses observed in previous AML trials.

KL50 provides a mechanistically distinct approach to exploiting the same MGMT vulnerability. Unlike TMZ-mediated methylation, KL50 deposits O^6^-fluoroethylguanine lesions that slowly evolve into DNA interstrand crosslinks (ICLs) ^35,43^. Our findings, together with previous work in solid tumor models ^22^, demonstrate that the processing and cytotoxicity of these lesions do not require functional MMR. Instead, KL50 shifts the repair burden toward replication-associated pathways, particularly Fanconi anemia (FA) and homologous recombination (HR) ^44^. This mechanistic distinction was reflected in our CRISPR screen and genetic validation studies, where loss of MMR conferred profound resistance to TMZ but provided no comparable fitness advantage under KL50 treatment. Consequently, KL50 retained activity in both intrinsically MMR-deficient AML and populations that acquired MMR-associated resistance following prolonged TMZ exposure. These findings distinguish O^6^-fluoroethylation from conventional O^6^-methylation and provide a strategy to exploit MGMT deficiency without inheriting one of the principal resistance mechanisms associated with TMZ. Our functional genomic analyses also identified additional modifiers of response. Loss of TP53 and CHEK2 emerged as prominent resistance-associated hits under KL50 selection, while FA and HR pathway genes represented major sensitizing dependencies. TP53 is particularly relevant given its association with treatment resistance and poor outcomes in AML ^45,46^. Importantly, KL50 alone and in combination with ATR inhibition remained active across genetically distinct models, including TP53-wild-type MOLM13 and TP53-deficient U937 and K562 backgrounds ^47,48,49,50^. Nevertheless, established cell lines cannot fully model the clonal complexity of TP53-mutant AML, and dedicated evaluation in genetically defined primary AML and PDX models will be important to determine how specific TP53 alterations influence KL50 response. More broadly, our focused 353-gene DDR CRISPR screen provides an initial map of the repair pathways governing response to O6-directed alkylation. Future genome-wide screening could extend this framework to identify non-canonical resistance mechanisms and additional synthetic-lethal dependencies beyond classical DNA repair pathways.

A critical requirement for translating this strategy is accurate identification of the MGMT-deficient population. MGMT promoter methylation has long served as a predictive biomarker for TMZ response in glioblastoma ^6,51,52^; however, promoter methylation does not invariably correspond to loss of functional MGMT protein ^29,53^. Our integrated analysis addresses this limitation by identifying a defined subset of MGMT promoter CpG sites whose methylation strongly associates with reduced MGMT transcript and protein abundance in AML. Importantly, healthy CD34^+^ hematopoietic stem and progenitor cells maintained substantially higher MGMT protein levels than MGMT-silenced AML samples. This differential expression provides a potential therapeutic window in which normal hematopoietic progenitors retain MGMT-mediated protection while MGMT-deficient leukemic cells remain selectively vulnerable to O^6^-directed damage. These findings support incorporating both optimized promoter methylation signatures and quantitative MGMT expression into future biomarker strategies rather than relying on broad methylation status alone.

Mechanistically, KL50-induced ICL formation generated pronounced replication stress, characterized by RPA32 phosphorylation and activation of checkpoint signaling. Although both ATM-CHK2 and ATR-CHK1 signaling were engaged, our combinatorial studies identified ATR inhibition as the strongest therapeutic interaction with KL50. This dependency provides a particularly attractive opportunity because ATR serves as a central survival pathway during replication stress, allowing cells to stabilize stalled replication forks and tolerate persistent DNA damage. Pharmacologic ATR inhibition therefore removes a critical survival mechanism precisely when MGMT-deficient AML cells are processing KL50-induced lesions. The clinical development of ATR inhibitors, however, has been constrained in part by hematologic toxicity resulting from inhibition of replication-stress responses in proliferating normal cells ^25,26^. We therefore used the known kinetics of KL50-induced lesion maturation to design a temporally sequenced combination strategy. Because O^6^-fluoroethylguanine lesions mature into ICLs slowly ^22,35^, delaying ATR inhibition for 24 hours allows KL50-induced lesions to develop before disrupting the subsequent ATR-dependent stress response. Sequential KL50 followed by Ceralasertib enhanced synergy *in vitro* and produced marked tumor growth inhibition and survival benefit across AML xenograft models without significant body-weight loss. Importantly, this interaction remained MGMT-dependent and MMR-independent, preserving the biomarker selectivity of KL50 while bypassing the resistance mechanism that limits TMZ.

Several translational questions remain. Although our studies establish efficacy across genetically diverse cell-line and primary AML models, long-term evaluation of the KL50-ATR inhibitor combination in additional PDX models will be important to assess efficacy, hematologic tolerability, and clonal evolution under sustained treatment pressure. Likewise, the activity of KL50 in TP53-mutant AML requires dedicated investigation given the resistance signal observed in our functional genomic screen. Finally, while the sequential regimen was well tolerated in our preclinical models, defining whether temporal separation can meaningfully reduce ATR inhibitor-associated myelosuppression will require detailed hematologic and bone marrow toxicity studies followed by clinical dose optimization

Collectively, our study establishes MGMT silencing as a prevalent therapeutic vulnerability in AML while identifying MMR deficiency as an important limitation of conventional O^6^-methylating therapy. By replacing methyl deposition with time-dependent O^6^-fluoroethylation and interstrand crosslink formation, KL50 preserves MGMT selectivity while bypassing MMR-dependent resistance and acquired clonal escape. The resulting replication stress creates an actionable ATR dependency that can be further exploited through rational temporal sequencing. Together, these findings provide a biomarker-driven framework for developing O^6^-fluoroethylating imidazotetrazines, alone or in combination with ATR inhibition, for de novo and R/R MGMT-silenced AML.

## Materials and Methods

### Chemicals

The O^6^-fluoroethylating triazene KL-50 was synthesized at Yale University or custom-synthesized by WuXi AppTec as previously described (1, 2). Temozolomide (TMZ), lomustine (CCNU), carmustine (BCNU), bendamustine, busulfan, ceralasertib (AZD6738), KU60019, prexasertib, and olaparib were purchased from Selleck Chemicals. Mitozolomide (MTZ) was purchased from Enamine, cyclophosphamide from Cayman Chemical, and methyl methanesulfonate (MMS) from Sigma-Aldrich.All compounds were dissolved in anhydrous DMSO to prepare master stocks at the following concentrations: 100 mM (KL50), 150 mM (TMZ, MTZ, busulfan), 100 mM (MMS, CCNU, BCNU, bendamustine), and 20 mM (cyclophosphamide). Master stocks were aliquoted and stored at −20 °C or −80 °C.

### ATCC cell lines and Primary patient samples

MOLM-13, U937, THP1, and K562 cell lines were cultured in RPMI 1640 (Gibco) supplemented with 10% fetal bovine serum (FBS; Sigma-Aldrich). Kasumi1 and HL60 cells were maintained in ATCC-modified RPMI and Iscove’s Modified Dulbecco’s Medium (IMDM; Gibco), respectively, each supplemented with 20% FBS. All cell lines were obtained from the ATCC, maintained at 37 °C under 5% CO_2_, and routinely authenticated via short tandem repeat (STR) profiling and mycoplasma testing (MycoAlert, Lonza).

Primary peripheral blood and bone marrow samples were procured from the Yale Hematology Tissue Bank under protocols approved by the Yale University Human Investigation Committee, following written informed consent from all donors. Mononuclear cells were isolated using Ficoll-Paque density gradient centrifugation (GE Healthcare) and cryopreserved in FBS containing 10% DMSO within 24 hours of collection.

Primary AML cells were cultured in DMEM (Gibco) supplemented with 10% FBS and a cytokine cocktail containing 50 ng/mL each of human recombinant stem cell factor (SCF), Flt3-ligand, and thrombopoietin (TPO), alongside 10 ng/mL IL-3 and 20 ng/mL IL-6. Healthy donor CD34+ cells were maintained in the identical cytokine-supplemented medium, excluding IL-6. All recombinant cytokines were sourced from Gemini Bio.

### Cell Line Engineering and Lentiviral Transduction

To generate MGMT isogenic models, MGMT was overexpressed in MOLM13, U937, K562, and HL60 cells using pGen-lenti-Neo-MGMT lentiviral vectors (GenScript), and knocked down in THP1 and Kasumi1 cells using custom MGMT-targeting shRNA pLKO.1-puro-CMV-tGFP lentiviral particles (Sigma-Aldrich). Mismatch repair (MMR)-deficient isogenic pairs were generated by transducing MOLM-13, U937, and K562 cells with MSH6-targeting pGIPZ shRNA lentiviral particles (Horizon Discovery). For *in vivo* bioluminescence tracking, MOLM-13 and U937 cells were stably transduced with CMV-Firefly Luciferase lentiviruses (Cellomics Technology). For all transductions, 2 million cells/mL were incubated with the respective lentiviral particles at a multiplicity of infection (MOI) of 0.5 in the presence of 8 µg/mL polybrene (Santa Cruz Biotechnology). Media was replenished after 24 hours to remove viral particles. Following an additional 48 hours of recovery, stable transductants were isolated via 5–7 days of antibiotic selection using either 2 mg/mL G418 (Sigma-Aldrich; for MGMT overexpression), 1 µg/mL puromycin (InvivoGen; for MGMT knockdown, and MSH6 knockdown), or 6 µg/mL blasticidin (Gibco; for luciferase expression).

### High-Throughput Cell Line Viability Screening (PRISM)

Multiplexed cellular viability screening was performed using the Profiling Relative Inhibition Simultaneously in Mixtures (PRISM) platform as previously described (Corsello et al., 2020 ^54^). KL-50 and TMZ were screened in triplicate across a molecularly barcoded library of approximately 900 cancer cell lines using an 8-point, 3-fold serial dilution profile for a 5-day treatment duration. The maximum assay concentration was 200 µM for KL-50 and 67 µM for TMZ, with the latter restricted due to solubility limitations at 200 µM. Bortezomib and DMSO served as positive and negative controls, respectively. Quality control filtering excluded cell lines with fewer than two passing replicates based on an error rate > 0.05 or a dynamic range < -log_2_(0.3), yielding 881 unique cell lines for downstream profiling. Viability metrics were calculated as normalized log-fold change values relative to DMSO vehicle controls. Dose-response curves were modeled using a four-parameter logistic fit to calculate absolute area under the curve (AUC) and IC50 values for each cell line model. Univariate associations between baseline MGMT expression and compound viability (AUC) were evaluated using Pearson correlation coefficients, with p-values adjusted for multiple hypothesis testing using the Benjamini–Hochberg false discovery rate algorithm (q-values). Baseline MGMT log2-transformed mRNA expression data and cell line metadata were extracted from the DepMap Portal Public 23Q4 release.

### Short-Term Cell Viability and Drug Synergy Assays

For baseline viability screenings, cells were seeded into the inner 48 wells of 96-well plates at a density of 1,000 cells/well in a final volume of 200 µl/well. Cells were treated with an 8-point, 2- or 3-fold serial dilution of the respective compounds. The final vehicle concentration was maintained at 0.5% DMSO across all treatment conditions, including the negative control wells. Following a 6-day incubation protocol, cell suspensions were transferred to white opaque 96-well microplates (Corning Costar) and mixed with CellTiter-Glo 2.0 reagent (Promega) at a 1:1 ratio. Plates were agitated on an orbital shaker for 2 minutes and incubated for 10 minutes at room temperature prior to luminescence acquisition on a BioTek Synergy H1 microplate reader. IC50 values were calculated using non-linear regression modeling (Inhibitor vs. normalized response with variable slope). For assays evaluating pharmacological MGMT depletion, cells were pretreated with 10 µM O^6^-benzylguanine (O6BG) for 1 hour prior to the addition of TMZ or KL-50; O6BG remained in the culture medium for the duration of the assay.

For checkerboard drug synergy assays, a similar experimental design was implemented with a final vehicle concentration of 1% DMSO. Cells were treated with an 8 x 6 matrix combining an 8-point serial dilution of the primary compound with a 6-point serial dilution of the secondary agent. Following a 6-day incubation, cellular ATP levels were quantified via CellTiter-Glo 2.0 as described above. Drug synergy was calculated based on HSA model using Combenefit software (Di Veroli et al., 2016 ^55^).

### Pooled CRISPR-Cas9 Knockout Screen and Analysis

MOLM13 cells were engineered to stably express Cas9 via transduction with the CRISPR Cas9 pR-CMV-Cas9-2A-Blast lentiviral construct (Cellecta) at a multiplicity of infection (MOI) of 0.25. The viral supernatant was replaced with fresh culture medium 24 hours post-infection. Following an additional 48 hours of recovery, stable Cas9-expressing pools were selected via sequential incubation with 6 µg/mL blasticidin (Gibco) for 14 days, followed by 24 µg/mL blasticidin for 7 days. To prevent single-clone artifacts, the total blasticidin-selected MOLM-13 population was retained as a pooled bulk culture. Efficacious Cas9 expression was confirmed via Western blot analysis utilizing an anti-Cas9 antibody (Cell Signaling Technology). A customized, pooled sgRNA library targeting 353 core DNA damage response (DDR) genes was designed and compiled as previously detailed (Heer et al., 2026 ^44^). Lentiviral screening protocols were adapted from Cellecta specifications and established laboratory workflows (Heer et al., 2026). Briefly, Cas9-expressing MOLM13 cells were transduced with the packaged sgRNA library at an MOI of 0.1–0.2 in the presence of 8 µg/mL polybrene (Sigma-Aldrich) and seeded into 15 cm dishes (Corning). Medium was replaced 24 hours post-transduction, and after 48 additional hours infected cells were selected with 800 µg/ml hygromycin (Invitrogen) for 5 days. Following selection (designated as Time 0), the actual screen MOI was verified, and the population was split into control and treatment arms. Cells were maintained at a minimum of 1,000-fold library coverage per replicate and treated in duplicate with either vehicle control (DMSO), temozolomide (TMZ), or KL-50 at pre-determined IC30 and IC87 screening concentrations. Cells were expanded under selective pressure, replated at 1,000-fold library coverage after 6 replication cycles (approximately 6-8 days), and retreated. After a cumulative 12 replication cycles (12–16 days), cell pellets comprising 4.5 million cells per condition were harvested, washed with PBS, flash-frozen, and stored at −80 °C. Genomic DNA isolation, guide RNA cassette amplification for high-throughput sequencing, quality control analysis, and MAGeCK algorithmic processing were executed as previously detailed (Heer et al., 2026). Functional classification and structural assignment of targeted genes into defined DDR pathways are cataloged in Supplementary Table 2.

### Western Blot Analysis and Downstream DDR Signaling/Apoptosis Assays

To evaluate baseline protein levels, cell pellets were lysed in RIPA buffer (50 mM HEPES, 250 mM NaCl, 5 mM EDTA, 1% NP-40) supplemented with 1X cOmplete EDTA-free protease inhibitor cocktail (Sigma-Aldrich) and PhosSTOP phosphatase inhibitor cocktail (Roche). Suspensions were sonicated (10 s on/10 s off for 1 min total) and cleared via centrifugation (13,000 rpm for 10 min at 4 °C). Protein concentrations were determined using Bradford, and samples were resolved on NuPAGE 4%–12% Bis-Tris gels (Thermo Fisher Scientific) before transfer onto nitrocellulose membranes (Sigma-Aldrich). Membranes were blocked for 1 hour at room temperature in either 5% bovine serum albumin (BSA) or 5% non-fat dry milk in Tris-buffered saline with 0.1% Tween-20 (TBST), according to target validation protocols. For baseline immunoblots, 20 µg of protein was loaded per lane. Primary antibodies targeting MGMT, MMR machinery, and housekeeping controls were incubated overnight at 4 °C at a 1:1,000 dilution in 5% BSA, with the following exceptions: anti-PMS2 was incubated for 1.5 hours at room temperature at 1:2,000, and anti-GAPDH was incubated at 1:5,000. Horseradish peroxidase (HRP)-conjugated secondary antibodies were applied at a 1:10,000 dilution for 1 hour at room temperature.

For replication stress, DNA damage response (DDR) signaling (pCHK1/2), and apoptosis kinetic assays, 10 million cells were seeded at a density of 1million cells/mL in 10 cm dishes and treated with 72 µM TMZ or KL-50 for 48 hours. Cells were harvested and lysed as detailed above, and 70 µg of total protein was loaded per lane. Primary antibodies tracking replication stress (p-RPA32 S33), DDR activation (p-CHK1, p-CHK2), and apoptosis induction (cleaved PAPR1, cleaved Caspases) were diluted 1:1,000 in 5% BSA or 5% milk (for phospho-proteins) and incubated overnight at 4 °C, except p-RPA32, which was diluted 1:10,000. Phospho-specific secondary antibodies were incubated for 2 hours at room temperature at either a 1:10,000 dilution in 5% milk or a 1:5,000 dilution (for p-RPA32 and p-CHK1). All blots were developed using Clarity or Clarity Max Western ECL Substrates (Bio-Rad) and visualized on Bio-Rad ChemiDoc or ChemiDoc Go imaging platforms. Comprehensive antibody parameters and catalog numbers are detailed in Supplementary Table 3.

### Gamma-H2AX Analysis

MOLM-13 and U937 cells were seeded in 6-well plates at a density of 2 million cells/well and immediately treated with TMZ or KL-50 for 48 hours (100 µM for MOLM-13; 20 µM for U937). Following incubation, cells were harvested via centrifugation (400 g for 3 min), washed with 1X PBS, and transferred to 96-well V-bottom plates. Cells were fixed with 4% paraformaldehyde for 30 minutes at room temperature, centrifuged, and permeabilized with 90% ice-cold methanol for 10 minutes at −20 °C. Following two successive washes with 1X PBS, cells were incubated overnight at 4 °C with an Alexa Fluor 647-conjugated anti-gamma-H2AX primary antibody (BioLegend #613407; 1:500 dilution in cell staining buffer). The cells were then centrifuged and treated with 100 µg/mL RNAse A (Qiagen) for 30 minutes, followed by a 1-hour incubation with Hoechst 33342 (20 µg/mL}; 1:50 dilution of a 1 mg/mL stock) for DNA content staining. After two final washes with 1X PBS, a minimum of 10,000 single-cell events were acquired on a CytoFLEX flow cytometer (Beckman Coulter). The percentage of gamma-H2AX-positive cells and their corresponding cell cycle phase distributions were quantified and analyzed using FlowJo software (version 10.10.0).

### Generation of Temozolomide-Resistant AML Sublines

To model acquired resistance, MGMT-deficient and mismatch repair (MMR)-proficient MOLM13 and U937 cells were subjected to chronic, long-term selection with temozolomide (TMZ). Cultures were treated weekly with step-wise increasing concentrations of TMZ over a duration of 14-15 weeks. Initial versus final selection thresholds were escalated as follows: 12 µM to 400 µM for MOLM-13, and 4 µM to 100 µM for U937.The emergence of stable resistance phenotypes was functionally validated using short-term CellTiter-Glo 2.0 viability assays. Concurrently, cell pellets were harvested from resistant sublines for downstream molecular profiling: total protein was extracted for Western blot validation and genomic DNA was isolated to characterize mutational signatures via Whole Exome Sequencing (WES).

### Animal studies

All animal experiments were executed under protocols approved by the Yale Institutional Animal Care and Use Committee (IACUC) and in accordance with the Association for Assessment and Accreditation of Laboratory Animal Care (AAALAC) guidelines.

#### In vivo Cell-derived xenotransplantation (CDX) studies

Athymic nude (Hsd:Athymic Nude-Foxn1nu) and NSG (NOD-scid IL2Rgammanull) mice were purchased from Envigo. Mice were housed up to five per individually ventilated cage at 20–24 °C and 40%–70% humidity, with access to food and water ad libitum. Animals were euthanized upon reaching humane endpoints, defined as severe morbidity, >20% body weight loss, or tumor volumes exceeding 2,000-3,000 mm^3^. For subcutaneous cell-derived xenograft (CDX) models, 2 million MOLM-13 cells were suspended in a 1:1 mixture of PBS and Matrigel (100 µl final volume) and injected into the flanks of 6- to 8-week-old athymic nude mice. Once tumors reached approximately 100 mm^3^, mice were randomized into treatment arms. Vehicle (10% cyclodextrin), KL-50 (5 mg/kg), or TMZ (25 mg/kg) were administered via oral gavage once daily on a 5-day on/2-day off schedule for 4–5 weeks. For drug combination studies, mice were randomized into four distinct arms: vehicle (10% cyclodextrin), KL-50 (5 mg/kg), AZD6738 (50 mg/kg), or a sequential combination of KL-50 (5 mg/kg) and AZD6738 (50 mg/kg). All compounds in the combination cohort were administered via oral gavage on a staggered, 24-hour sequential dosing schedule (KL-50 on Days 1 and 4; AZD6738 on Days 2 and 5 of a weekly cycle) for 3-4 weeks. Tumor dimensions were monitored twice weekly using electronic calipers, and volumes were calculated using the formula V = 0.523 x Length x Width^2^. Animal survival kinetics were modeled via Kaplan-Meier analysis.

For intravenously (i.v.) engrafted AML models, 6- to 8-week-old NSG mice were pre-conditioned with 2.25 Gy of total body irradiation using an X-Rad 320 X-ray irradiator. Approximately 24 hours post-irradiation, 2 million luciferase-expressing U937 cells were suspended in 100 µl of PBS and injected via the lateral tail vein. Baseline bioluminescence imaging (BLI; designated as Time 0) was performed immediately following engraftment by administering D-luciferin (150 mg/kg) via intraperitoneal injection 10 minutes prior to image acquisition on an IVIS Spectrum In Vivo Imaging System (PerkinElmer). Mice were randomized based on their Time 0 BLI signal intensity, and therapies were initiated 5 days post-injection to allow stable engraftment. Follow-up BLI was performed on Day 5 and tracked once weekly thereafter. For both monotherapy and combination arms, the compound dosages and sequential scheduling mirrored the protocols outlined for the subcutaneous models. Total tumor burden was monitored by quantifying the average bioluminescent flux (photons/second/cm^2^) using the entire mouse body as the region of interest (ROI). Animals were monitored continuously to evaluate survival kinetics until reaching humane endpoints.

#### Development of TMZ-induced CDX models

To develop *in vivo* models of acquired TMZ resistance, subcutaneous MOLM-13 tumors were established in athymic nude mice as detailed above. Upon reaching 100 mm^3^, mice were randomized and treated chronically with 25 mg/kg TMZ via oral gavage (5 days on/2-day off weekly schedule). Treatment was maintained continuously until the tumors escaped selective pressure and progressed to the maximum humane endpoint. Refractory tumors were then harvested and enzymatically dissociated into single-cell suspensions. These acquired-resistance sublines were expanded in vitro for downstream molecular and genomic validation using Western blotting and whole-exome sequencing (WES)

#### Engraftment and Analysis of MGMT-Stratified AML Patient Samples

MISTRG6 mice (CSF1h/h IL3/CSF2h/h SIRPAh/m THPOh/h Rag2–/– Il2rg–/– IL6h/h) were generated by crossing MISh/hTRG6 and MISm/mTRG6 strains to produce human cytokine homozygous and human SIRPA (hSIRPA) heterozygous offspring ^56^. These mice (referred to as MISTRG6 throughout the study) have been deposited at The Jackson Laboratory; requests for access via Material Transfer Agreement (MTA) should be directed to.

For transplantation, AML patient samples, stratified as MGMT-high and MGMT-low, were injected into 3-day-old newborn MISTRG6 mice following a single 200 cGy dose of irradiation (XRAD 320, PXI X-Ray Systems). Cryopreserved BM and splenocytes derived from patient-derived xenografts (PDXs) were thawed and incubated with an anti-human CD3 antibody (clone OKT3, BioXCell, Cat# BE0001-2, RRID: AB_1107632) at 5 µg/100 µl for 10 minutes at room temperature to prevent Graft-versus-Host Disease (GvHD). Cells were then delivered intra-hepatically in a 20µl volume using a 22-gauge Hamilton needle and syringe (Hamilton, Reno). Between 6 and 16 weeks post-transplantation, PB was collected for analysis. Hgb and platelets were measured via CBC, while human CD45+ (huCD45+) and human CD34+ (huCD34+) engraftment levels were quantified by flow cytometry. Mice were then assigned to balanced treatment groups (Vehicle, KL50, or TMZ). Vehicle (10% cyclodextrin), KL50 (5 mg/kg), and Temozolomide (TMZ) (25 mg/kg) were administered via oral gavage once daily for four weeks (5 days on / 2 days off per week). Following the initiation of treatment, mice were weighed twice weekly and monitored every two weeks via complete blood counts (CBC) and flow cytometry to track engraftment levels (Supplementary Figure S4B). The Y2315 and Y4008 cohorts were followed for total durations of 180 and 150 days, respectively. Mice were harvested either upon reaching humane endpoints due to deteriorating health or at the scheduled experiment termination. For the engraftment analysis of human CD45^+^ cells and their subset, cells were isolated from engrafted mice, blocked with human BD Fc block antibody (STEMCELL Technologies, IV.3, Cat#60012, RRID:AB_2925215) and mouse BD Fc block (2.4G2, BD Pharmingen, Cat#553141, RRID: AB_394656), and stained with a combination of antibodies purchased from Biolegend: Human cell engraftment panel: APC/Cy7 mCD45 (clone 30-F11, 1:200, Cat#103116, RRID: AB_312981), APC/Cy7 mTer119 (clone TER-119, 1:200, Cat#116223, RRID: AB_2137788), BV510 huCD45 (clone HI30, 1:100, Cat#304036, RRID: AB_2561940), FITC huCD3 (clone UCHT1, 1:100, Cat#300406, RRID: AB_314060), PE/Cy7 huCD19 (clone HIB19,1:100, Cat#302216, RRID: AB_314246), APC huCD33 (clone WM53, 1:100, Cat#303408, AB_314352), PE huCD34 (clone 561, 1:100, Cat#343606, AB_1732008). Spectral flow cytometry was performed on a Cytek Aurora (5-laser, 64 detector) using SpectroFlo software (Cytek Biosciences) at the Yale Flow Cytometry Core Facility and analyzed with FlowJo v10.10 software (BD Biosciences, Ashland, OR).

### RNA-seq protocol and analysis

#### TCGA RNA-seq data analysis for AML and pan-cancer

Data Sources: TCGA RNA-seq. Per-sample gene-level read counts and TPM (tpm_unstranded) from the GDC STAR – Counts workflow ^57^ were downloaded with gdc-client using manifest gdc_manifest.2025-10-23.112354.txt (10,048 primary-tumor RNA-seq files spanning 33 TCGA projects).

Clinical metadata: Tumor-type acronyms came from clinical_PANCAN_patient_with_followup.tsv and TCGA-CDR-SupplementalTableS1.xlsx ^58^. All TCGA samples were keyed on the sample barcode (TCGA-XX-XXXX-XXX).

Per-sample GDC STAR-counts files (one *.rna_seq.augmented_star_gene_counts.tsv per file in the manifest above) were parsed and concatenated into a long-format expression table indexed on the sample barcode and gene symbol. MGMT TPM was extracted across all TCGA-LAML samples (AML cohort) and across all 33 TCGA projects (pan-cancer cohort); samples without a usable count file were excluded.

#### Yale RNA-seq data analysis

RNA from AML patient bone marrow cells (Yale Hematology Biobank) was isolated using RNeasy kit (Qiagen) and RNA-seq (poly A) was performed on Novaseq at the Yale Centre for Genomic Analysis core facility. Paired-end FASTQ files from Yale patient samples were processed with the nf-core/rnaseq Nextflow pipeline (v3.12.0) ^59^ in star_salmon mode against the Ensembl GRCh38 primary assembly with annotation Ensembl release 112 ^60^. Reads were adapter-trimmed with Trim Galore v0.6.7 / Cutadapt v3.4 [Krueger 2023; Martin 2011], aligned with STAR v2.7.10a ^61^, and quantified with Salmon v1.10.1 ^62^. Gene-level TPM was used for downstream visualizations.

### Methylation array protocol and analysis

#### Probe-level correlation of methylation with MGMT mRNA expression in TCGA AML and pan-cancer datasets

Data source: Two complementary β-value resources were used. A) Broad FireHose Level 3 HumanMethylation450 matrices for LAML (stddata__2016_01_28 release [Broad 2016]), retaining only the β-value channel from the four-column-per-sample format. B) TCGA PanCancer Methylation450 whitelisted β-value matrix jhu-usc.edu_PANCAN_HumanMethylation450.betaValue_whitelisted.tsv ^63^. For samples present in both resources, FireHose values were preferred during de-duplication.

First, cross-platform methylation probe harmonization was done. CpG probe coordinates from the Illumina HumanMethylation27, HumanMethylation450, and Infinium MethylationEPIC v2.0 (8v2-0_A1) manifests [Illumina 2008, 2011, 2024] were unified to GRCh38 by liftOver ^64^ or used directly (EPIC v2). Where multiple manifest entries described the same IlmnID, the entry with the fewest missing annotation fields was retained. The harmonized panel was filtered to probes mapping within ± 100 bp of the MGMT gene body plus any probe in the per-disease correlated-probe sets described in section 2. EPIC v2 probes sharing an underlying CpG (collapsed at the IlmnID prefix) were averaged per sample. This procedure was applied independently to each cohort (LAML, and pan-cancer). HumanMethylation450 β-values from the TCGA PanCancer whitelisted matrix and gene-level expression level from the legacy TCGA PanCancer batch-corrected expression log2(RSEM + 1) matrix EBPlusPlusAdjustPANCAN_IlluminaHiSeq_RNASeqV2.geneExp.tsv ^63^ were aggregated to the sample level (mean across replicate samples) and merged on the sample barcode. Probes were retained if they overlapped the gene-of-interest locus (± 100 bp) on hg19 or were annotated to that gene in the HM450 UCSC_RefGene_Name field. Per-probe Pearson correlations between β-value and expression, two-sided p-value, and probe coordinate were computed. The eleven MGMT-region probes most strongly negatively correlated with MGMT expression in the pan-cancer / AML cohorts were carried forward to downstream analysis.

#### Yale methylation analysis (Illumina EPIC v2 array)

Genomic DNA from AML patient bone marrow cells (Yale Hematology Biobank) was isolated using DNeasy blood and tissue kit (Qiagen). Genomic DNA was bisulfite-converted and analyzed for genome-wide methylation patterns using the Illumina Infinium MethylationEPIC v2.0 BeadChip (8v2-0_A1) [Illumina 2024] at the Keck Microarray core facility at Yale. Raw IDAT files were processed in Illumina GenomeStudio (Methylation Module) using control-probe normalization to obtain per-sample AVG_Beta and total-intensity values [Illumina 2020]. Probes with any null AVG_Beta across the cohort were excluded; replicate measurements sharing an underlying CpG (EPIC v2 IlmnID prefix) were collapsed to the mean β.

#### Softwares

Data transformation and analysis was performed in Python 3.11 with pandas [McKinney 2010], NumPy ^65^, SciPy ^66^, and pyranges ^67^ and in R with minfi ^68^. Workflow orchestration used Nextflow ^69^ within Singularity containers.

### Whole-exome sequencing protocol and analysis

Genomic DNA from the samples was isolated using DNeasy blood and tissue kit (Qiagen) and submitted at >20 ng/µl concentration to the Yale Centre for Genomic Analysis core facility for whole exome sequencing. Regarding the analysis, the pipeline used followed the GATK 4 best practices guidelines for generating called variants ^70^. The analysis pipeline aligns the paired-end reads using BWA MEM ^71^ to the hs38DH human reference, which are versions of the hg38 reference that includes decoy sequences. It then marks PCR duplicates using picard’s MarkDuplicates command. The GATK 4 software ^72^ is used to recalibrate base quality scores and to generate CRAM and GVCF files for each sample. Once GVCF files have been generated, joint variant calling is performed and variants are filtered using either hard filtering (if a small number of samples are involved) or variant quality score recalibration (if a whole cohort is being analyzed). All of the samples, across all groups, were joint-called together.

VCF files are annotated using the ANNOVAR ^73^ and Variant Effect Predictor (VEP) ^74^, as well as additional databases such as OMIM (https://www.omim.org/), GO (http://geneontology.org/) and ClinVar (https://www.ncbi.nlm.nih.gov/clinvar/), in order to capture (1) the prediction of the variants’ functional effects (damaging nonsense, splice, missense protein altering. or frameshift variants), (2) whether the variants are common or rare in the population, (3) the conservation/intolerance of the variant or the gene across related species and (4) any relationships between the genes/variants known phenotypes and the disease phenotype.

The annotated variants were then filtered for variants that are enriched in the treated samples, compared to the group’s wildtype sample. The initial filtering kept medium and high impact variants (those that change the transcribed protein), and then a data-driven, filtering refinement process was performed, as described next. The filtering criteria from that process was used for each Treated-WT pair to generate the final set of variants.

The objective of this analysis is to identify any variant whose allele frequency changes as a result of the treatment, as compared to the wildtype, while filtering out sequencing errors. This makes it more general than the “de novo variant” method used in somatic variant callers. To achieve this, a “reciprocal subtraction and refinement” method was used on the filtering criteria. In each round of the refinement, the current filtering criteria were used on each Treated-WT pair, to generate variants that are Treated-enriched (by subtracting variants found in WT) and WT-enriched (by subtracting Treated variants).

The expectation is that the WT-enriched variants are a result of sequencing errors not caught by the filtering criteria, and those were evaluated to identify additional sequencing error modes that should be filtered. After several round of refinements, the final set of filtering criteria were the following (where Ref here refers to the sample used for subtraction and Query refers to the “enriched” sample): i) All variants for the pair were also called by the Freebayes variant caller, using its “force calling” method with options “-t variants.bed -l -@ variants.vcf”, where variants.vcf is a vcf file of the pair’s variants, and variants.bed is a bed file listing the loci of the variants. ii) Variants not called by both variant callers, GATK and Freebayes, were filtered. iii) Variants where “abs(GATK Query AF – Freebayes Query AF) > 0.15” were filtered (i.e., where GATK and Freebayes Query AF differed by more than 15%). iv) Variants where, for GATK, “abs(Ref AF – Query AF) < 0.15” were filtered (i.e., the Ref and Query AF differed by less than 15%). v) Filtered variants where “Freebayes Ref GQ < 50” or “Freebayes Query GQ < 80”. vi) Filtered variants where the Freebayes mean basecall quality score (using all reads) was less than 26.

One key reason for the use of Freebayes (apart from having a second caller to filter out miscalls from GATK) is that the NovaSeq XPlus has a new error mode, where sequencing errors with high allele frequencies (up to 30-40%) are only identified by the low basecall quality of the reads (in this error mode, both ref and alt reads are called with low-quality scores). The GATK basecall quality metric expects high basecall quality for ref calls but low quality for the sequencing error calls, and so does not identify this mode where the ref calls also have low basecall quality. Freebayes provides basecall quality metrics that allow for the mean basecall quality score to be computed.

For studying acquisition of damaging mutations in MMR genes, we computed paired pooled damaging-to-all VAF effects for MMR genes versus exome background across predefined baseline and treated sample comparisons: from annotated exome calls we retained ‘PASS’ variants with VAF>0 and labeled variants as damaging if CADD≥20 or truncating (frameshift, stop_gained, splice_acceptor/donor); per sample we aggregated per⍰gene SumVAF_All and SumVAF_Damaging and formed per⍰gene R = SumVAF_Damaging / SumVAF_All. For each pair we pooled R across the union of observed MMR genes and across eligible background genes (non⍰MMR genes with Avg_SumVAF_All above the 25th⍰percentile mass floor) and computed the primary statistic Observed_DiD = (R_MMR_treated − R_MMR_baseline) − (R_BG_treated − R_BG_baseline), which was represented as Delta (Δ) damaging VAF fraction in the Supplementary Figures S6B,F.

### Statistical Analysis

All statistical analyses were performed using GraphPad Prism (version 10.6.0). Data distribution was first assessed using the Shapiro**-**Wilk normality test. For data following a normal distribution, comparisons between two groups were performed using an unpaired Student’s t-test when variances were not significantly different; otherwise, Welch’s unpaired t-test was applied. For non-normally distributed data, comparisons between two groups were made using the non-parametric Mann**-**Whitney test. For comparisons involving more than two groups, an ordinary one-way ANOVA with Dunnett’s multiple comparisons correction was used for normally distributed data, whereas the non-parametric Kruskal**-**Wallis test followed by Dunn’s multiple comparisons correction was employed for non-normally distributed data. Survival analyses were performed using the log-rank (Mantel**-**Cox) test applied to Kaplan**-**Meier survival curves. Statistical significance is indicated in all figures as follows: p < 0.05 (**\***), p < 0.01 (**\*\***), p < 0.001 (**\*\*\***), p < 0.0001 (****), and p > 0.05 (ns, not significant).

## Acknowledgements

We are deeply grateful to the patients who generously contributed primary samples for this research. Their participation and willingness to support scientific discovery made this work possible. We thank Yale Center for Genome Analysis (YCGA) and Keck Microarray Shared Resource Laboratory for their multi-omics (genomic, transcriptomic, and methylomics) support and data analysis. We would also like to specifically thank Dr. James Knight for whole-exome sequencing data analysis. We thank the Yale Hematology Tissue Bank and the DeLuca Center for Innovation in Hematology Research for provision of primary patient samples. We also thank the Irradiator Shared Resource by Yale Cancer Centre for small animal irradiation support and Yale Animal Resources Center for animal care. We thank Yale Flow Cytometry Core for flow cytometry analysis.

## Author Contributions

Conceptualization: PB, AB, SEG, SH, and RSB; Methodology: PB, AB, CDH, SEG, SH, RSB; Investigation: PB, AB, RKS, SF, JLE, AK, MM; Data analysis: PB, AB, SF, JLE, JV, CDH; Validation: PB, AB; Writing original draft: PB, AB; Writing, review & editing: PB, AB, SH, RSB; Funding acquisition: SH and RSB; Resources: SH and RSB; Project administration: RKS, SH, and RSB; Supervision: SH, and RSB.

## References

1. Siegel RL, Kratzer TB, Wagle NS, Sung H, Jemal A. Cancer statistics, 2026. CA Cancer J Clin. 2026;76(1):e70043.

2. Dohner H, Estey E, Grimwade D, et al. Diagnosis and management of AML in adults: 2017 ELN recommendations from an international expert panel. Blood. 2017;129(4):424–447.

3. Knijnenburg TA, Wang L, Zimmermann MT, et al. Genomic and Molecular Landscape of DNA Damage Repair Deficiency across The Cancer Genome Atlas. Cell Rep. 2018;23(1):239–254 e236.

4. Pegg AE. Mammalian O6-alkylguanine-DNA alkyltransferase: regulation and importance in response to alkylating carcinogenic and therapeutic agents. Cancer Res. 1990;50(19):6119–6129.

5. Thomas A, Tanaka M, Trepel J, Reinhold WC, Rajapakse VN, Pommier Y. Temozolomide in the Era of Precision Medicine. Cancer Res. 2017;77(4):823–826.

6. Esteller M, Toyota M, Sanchez-Cespedes M, et al. Inactivation of the DNA repair gene O6-methylguanine-DNA methyltransferase by promoter hypermethylation is associated with G to A mutations in K-ras in colorectal tumorigenesis. Cancer Res. 2000;60(9):2368–2371.

7. Aquilina G, Biondo R, Dogliotti E, Meuth M, Bignami M. Expression of the endogenous O6-methylguanine-DNA-methyltransferase protects Chinese hamster ovary cells from spontaneous G:C to A:T transitions. Cancer Res. 1992;52(23):6471–6475.

8. Hegi ME, Diserens AC, Gorlia T, et al. MGMT gene silencing and benefit from temozolomide in glioblastoma. N Engl J Med. 2005;352(10):997–1003.

9. Lu C, Ward PS, Kapoor GS, et al. IDH mutation impairs histone demethylation and results in a block to cell differentiation. Nature. 2012;483(7390):474–478.

10. Figueroa ME, Abdel-Wahab O, Lu C, et al. Leukemic IDH1 and IDH2 mutations result in a hypermethylation phenotype, disrupt TET2 function, and impair hematopoietic differentiation. Cancer Cell. 2010;18(6):553–567.

11. Turcan S, Rohle D, Goenka A, et al. IDH1 mutation is sufficient to establish the glioma hypermethylator phenotype. Nature. 2012;483(7390):479–483.

12. Medeiros BC, Kohrt HE, Gotlib J, et al. Tailored temozolomide therapy according to MGMT methylation status for elderly patients with acute myeloid leukemia. Am J Hematol. 2012;87(1):45–50.

13. Brandwein JM, Kassis J, Leber B, et al. Phase II study of targeted therapy with temozolomide in acute myeloid leukaemia and high-risk myelodysplastic syndrome patients pre-screened for low O(6) -methylguanine DNA methyltransferase expression. Br J Haematol. 2014;167(5):664–670.

14. Cheng X, An J, Lou J, et al. Trans-lesion synthesis and mismatch repair pathway crosstalk defines chemoresistance and hypermutation mechanisms in glioblastoma. Nat Commun. 2024;15(1):1957.

15. Touat M, Li YY, Boynton AN, et al. Mechanisms and therapeutic implications of hypermutation in gliomas. Nature. 2020;580(7804):517–523.

16. Yip S, Miao J, Cahill DP, et al. MSH6 mutations arise in glioblastomas during temozolomide therapy and mediate temozolomide resistance. Clin Cancer Res. 2009;15(14):4622–4629.

17. Crisafulli G, Sartore-Bianchi A, Lazzari L, et al. Temozolomide Treatment Alters Mismatch Repair and Boosts Mutational Burden in Tumor and Blood of Colorectal Cancer Patients. Cancer Discov. 2022;12(7):1656–1675.

18. Germano G, Lamba S, Rospo G, et al. Inactivation of DNA repair triggers neoantigen generation and impairs tumour growth. Nature. 2017;552(7683):116–120.

19. Wang J, Cazzato E, Ladewig E, et al. Clonal evolution of glioblastoma under therapy. Nat Genet. 2016;48(7):768–776.

20. Johnson BE, Mazor T, Hong C, et al. Mutational analysis reveals the origin and therapy-driven evolution of recurrent glioma. Science. 2014;343(6167):189–193.

21. Mao G, Yuan F, Absher K, et al. Preferential loss of mismatch repair function in refractory and relapsed acute myeloid leukemia: potential contribution to AML progression. Cell Res. 2008;18(2):281–289.

22. Lin K, Gueble SE, Sundaram RK, Huseman ED, Bindra RS, Herzon SB. Mechanism-based design of agents that selectively target drug-resistant glioma. Science. 2022;377(6605):502–511.

23. Hickman MJ, Samson LD. Role of DNA mismatch repair and p53 in signaling induction of apoptosis by alkylating agents. Proc Natl Acad Sci U S A. 1999;96(19):10764–10769.

24. Kaina B, Christmann M. DNA repair in personalized brain cancer therapy with temozolomide and nitrosoureas. DNA Repair (Amst). 2019;78:128–141.

25. Jurczak W, Elmusharaf N, Fox CP, et al. Phase I/II results of ceralasertib as monotherapy or in combination with acalabrutinib in high-risk relapsed/refractory chronic lymphocytic leukemia. Ther Adv Hematol. 2023;14:20406207231173489.

26. Barnieh FM, Loadman PM, Falconer RA. Progress towards a clinically-successful ATR inhibitor for cancer therapy. Curr Res Pharmacol Drug Discov. 2021;2:100017.

27. Shih AH, Abdel-Wahab O, Patel JP, Levine RL. The role of mutations in epigenetic regulators in myeloid malignancies. Nat Rev Cancer. 2012;12(9):599–612.

28. Abdel-Wahab O, Levine RL. Mutations in epigenetic modifiers in the pathogenesis and therapy of acute myeloid leukemia. Blood. 2013;121(18):3563–3572.

29. Zappe K, Puhringer K, Pflug S, et al. Association between MGMT Enhancer Methylation and MGMT Promoter Methylation, MGMT Protein Expression, and Overall Survival in Glioblastoma. Cells. 2023;12(12).

30. Cabrini G, Fabbri E, Lo Nigro C, Dechecchi MC, Gambari R. Regulation of expression of O6-methylguanine-DNA methyltransferase and the treatment of glioblastoma (Review). Int J Oncol. 2015;47(2):417–428.

31. Nakagawachi T, Soejima H, Urano T, et al. Silencing effect of CpG island hypermethylation and histone modifications on O6-methylguanine-DNA methyltransferase (MGMT) gene expression in human cancer. Oncogene. 2003;22(55):8835–8844.

32. Pegg AE. Multifaceted roles of alkyltransferase and related proteins in DNA repair, DNA damage, resistance to chemotherapy, and research tools. Chem Res Toxicol. 2011;24(5):618–639.

33. Woo PYM, Li Y, Chan AHY, et al. A multifaceted review of temozolomide resistance mechanisms in glioblastoma beyond O-6-methylguanine-DNA methyltransferase. Glioma. 2019;2(2):68–82.

34. Beranek DT. Distribution of methyl and ethyl adducts following alkylation with monofunctional alkylating agents. Mutat Res. 1990;231(1):11–30.

35. Huseman ED, Lo A, Fedorova O, et al. Mechanism of Action of KL-50, a Candidate Imidazotetrazine for the Treatment of Drug-Resistant Brain Cancers. J Am Chem Soc. 2024;146(27):18241–18252.

36. Jiricny J. The multifaceted mismatch-repair system. Nat Rev Mol Cell Biol. 2006;7(5):335–346.

37. Sarkaria JN, Kitange GJ, James CD, et al. Mechanisms of chemoresistance to alkylating agents in malignant glioma. Clin Cancer Res. 2008;14(10):2900–2908.

38. Fink D, Aebi S, Howell SB. The role of DNA mismatch repair in drug resistance. Clin Cancer Res. 1998;4(1):1–6.

39. Pu J, Yuan K, Tao J, et al. Glioblastoma multiforme: an updated overview of temozolomide resistance mechanisms and strategies to overcome resistance. Discov Oncol. 2025;16(1):731.

40. Dohner H, Wei AH, Lowenberg B. Towards precision medicine for AML. Nat Rev Clin Oncol. 2021;18(9):577–590.

41. Ding L, Ley TJ, Larson DE, et al. Clonal evolution in relapsed acute myeloid leukaemia revealed by whole-genome sequencing. Nature. 2012;481(7382):506–510.

42. van Galen P, Hovestadt V, Wadsworth Ii MH, et al. Single-Cell RNA-Seq Reveals AML Hierarchies Relevant to Disease Progression and Immunity. Cell. 2019;176(6):1265–1281 e1224.

43. Tong WP, Kirk MC, Ludlum DB. Mechanism of action of the nitrosoureas--V. Formation of O6-(2-fluoroethyl)guanine and its probable role in the crosslinking of deoxyribonucleic acid. Biochem Pharmacol. 1983;32(13):2011–2015.

44. Heer CD, Elia JL, Menon V, et al. Targeted CRISPR knockout screening identifies known and novel chemogenomic interactions between DNA damaging agents and DNA repair genes. NAR Cancer. 2026;8(1):zcaf052.

45. Shahzad M, Amin MK, Daver NG, et al. What have we learned about TP53-mutated acute myeloid leukemia? Blood Cancer J. 2024;14(1):202.

46. Granowicz EM, Jonas BA. Targeting TP53-Mutated Acute Myeloid Leukemia: Research and Clinical Developments. Onco Targets Ther. 2022;15:423–436.

47. Wang E, Pineda JMB, Kim WJ, et al. Modulation of RNA splicing enhances response to BCL2 inhibition in leukemia. Cancer Cell. 2023;41(1):164–180 e168.

48. Stubbs MC, Kim YM, Krivtsov AV, et al. MLL-AF9 and FLT3 cooperation in acute myelogenous leukemia: development of a model for rapid therapeutic assessment. Leukemia. 2008;22(1):66–77.

49. Sugimoto K, Toyoshima H, Sakai R, et al. Frequent mutations in the p53 gene in human myeloid leukemia cell lines. Blood. 1992;79(9):2378–2383.

50. Boettcher S, Miller PG, Sharma R, et al. A dominant-negative effect drives selection of TP53 missense mutations in myeloid malignancies. Science. 2019;365(6453):599–604.

51. Esteller M, Hamilton SR, Burger PC, Baylin SB, Herman JG. Inactivation of the DNA repair gene O6-methylguanine-DNA methyltransferase by promoter hypermethylation is a common event in primary human neoplasia. Cancer Res. 1999;59(4):793–797.

52. Zhang JY, Rajendran BK, Desai SS, et al. High MGMT expression identifies aggressive colorectal cancer with distinct genomic features and immune evasion properties. J Immunother Cancer. 2025;13(9).

53. Toffolatti L, Scquizzato E, Cavallin S, et al. MGMT promoter methylation and correlation with protein expression in primary central nervous system lymphoma. Virchows Arch. 2014;465(5):579–586.

54. Corsello SM, Nagari RT, Spangler RD, et al. Discovering the anti-cancer potential of non-oncology drugs by systematic viability profiling. Nat Cancer. 2020;1(2):235–248.

55. Di Veroli GY, Fornari C, Wang D, et al. Combenefit: an interactive platform for the analysis and visualization of drug combinations. Bioinformatics. 2016;32(18):2866–2868.

56. Das R, Strowig T, Verma R, et al. Microenvironment-dependent growth of preneoplastic and malignant plasma cells in humanized mice. Nat Med. 2016;22(11):1351–1357.

57. Heath AP, Ferretti V, Agrawal S, et al. The NCI Genomic Data Commons. Nat Genet. 2021;53(3):257–262.

58. Liu J, Lichtenberg T, Hoadley KA, et al. An Integrated TCGA Pan-Cancer Clinical Data Resource to Drive High-Quality Survival Outcome Analytics. Cell. 2018;173(2):400–416 e411.

59. Ewels PA, Peltzer A, Fillinger S, et al. The nf-core framework for community-curated bioinformatics pipelines. Nat Biotechnol. 2020;38(3):276–278.

60. Harrison PW, Amode MR, Austine-Orimoloye O, et al. Ensembl 2024. Nucleic Acids Res. 2024;52(D1):D891–D899.

61. Dobin A, Davis CA, Schlesinger F, et al. STAR: ultrafast universal RNA-seq aligner. Bioinformatics. 2013;29(1):15–21.

62. Patro R, Duggal G, Love MI, Irizarry RA, Kingsford C. Salmon provides fast and bias-aware quantification of transcript expression. Nat Methods. 2017;14(4):417–419.

63. Hoadley KA, Yau C, Hinoue T, et al. Cell-of-Origin Patterns Dominate the Molecular Classification of 10,000 Tumors from 33 Types of Cancer. Cell. 2018;173(2):291–304 e296.

64. Hinrichs AS, Karolchik D, Baertsch R, et al. The UCSC Genome Browser Database: update 2006. Nucleic Acids Res. 2006;34(Database issue):D590–598.

65. Harris CR, Millman KJ, van der Walt SJ, et al. Array programming with NumPy. Nature. 2020;585(7825):357–362.

66. Virtanen P, Gommers R, Oliphant TE, et al. SciPy 1.0: fundamental algorithms for scientific computing in Python. Nat Methods. 2020;17(3):261–272.

67. Stovner EB, Saetrom P. PyRanges: efficient comparison of genomic intervals in Python. Bioinformatics. 2020;36(3):918–919.

68. Aryee MJ, Jaffe AE, Corrada-Bravo H, et al. Minfi: a flexible and comprehensive Bioconductor package for the analysis of Infinium DNA methylation microarrays. Bioinformatics. 2014;30(10):1363–1369.

69. Di Tommaso P, Chatzou M, Floden EW, Barja PP, Palumbo E, Notredame C. Nextflow enables reproducible computational workflows. Nat Biotechnol. 2017;35(4):316–319.

70. Van der Auwera GA, Carneiro MO, Hartl C, et al. From FastQ data to high confidence variant calls: the Genome Analysis Toolkit best practices pipeline. Curr Protoc Bioinformatics. 2013;43(1110):11 10 11–11 10 33.

71. Li H, Durbin R. Fast and accurate short read alignment with Burrows-Wheeler transform. Bioinformatics. 2009;25(14):1754–1760.

72. McKenna A, Hanna M, Banks E, et al. The Genome Analysis Toolkit: a MapReduce framework for analyzing next-generation DNA sequencing data. Genome Res. 2010;20(9):1297–1303.

73. Wang K, Li M, Hakonarson H. ANNOVAR: functional annotation of genetic variants from high-throughput sequencing data. Nucleic Acids Res. 2010;38(16):e164.

74. McLaren W, Gil L, Hunt SE, et al. The Ensembl Variant Effect Predictor. Genome Biol. 2016;17(1):122.

## References

Broad Institute TCGA Genome Data Analysis Center. (2016). Standardized data run, January 28, 2016 (stddata__2016_01_28) [Data set]. Broad Institute of MIT and Harvard. 10.7908/C11G0KM9

Illumina, Inc. (2008). Infinium HumanMethylation27 BeadChip product support files [Manifest / content list]. https://support.illumina.com/downloads/humanmethylation27_product_support_files.html

Illumina, Inc. (2011). Infinium HumanMethylation450 BeadChip downloads (Manifest v1.2) [Manifest]. https://support.illumina.com/array/array_kits/infinium_humanmethylation450_beadchip_kit/downloads.html

Illumina, Inc. (2023). Infinium MethylationEPIC v2.0 Kit. https://www.illumina.com/products/by-type/microarray-kits/infinium-methylation-epic.html

Illumina, Inc. (2020). GenomeStudio software (Methylation Module 2011.1; framework v2.0.5) [Computer software]. https://support.illumina.com/array/array_software/genomestudio/downloads.html

McKinney, W. (2010). Data structures for statistical computing in Python. In S. van der Walt & J. Millman (Eds.), Proceedings of the 9th Python in Science Conference (pp. 56–61). 10.25080/Majora-92bf1922-00a

Waskom, M. L. (2021). seaborn: Statistical data visualization. Journal of Open Source Software, 6(60), 3021. 10.21105/joss.03021

Patel, H., Ewels, P., Peltzer, A., Hammarén, R., Botvinnik, O., Sturm, G., … nf-core community. (2023). nf-core/rnaseq: nf-core/rnaseq v3.12.0 [Computer software]. Zenodo. 10.5281/zenodo.1400710

Krueger, F. (2023). Trim Galore (v0.6.7) [Computer software]. Babraham Bioinformatics. https://github.com/FelixKrueger/TrimGalore

Martin, M. (2011). Cutadapt removes adapter sequences from high-throughput sequencing reads. EMBnet.journal, 17(1), 10–12. 10.14806/ej.17.1.200

