## Supplemental Figures 1-7 for "Biomarker-Targeted O^6^-Guanine Alkylation Potentiates ATR Inhibitor Response in De Novo and Relapsed/Refractory AML"

### Supplementary Figures

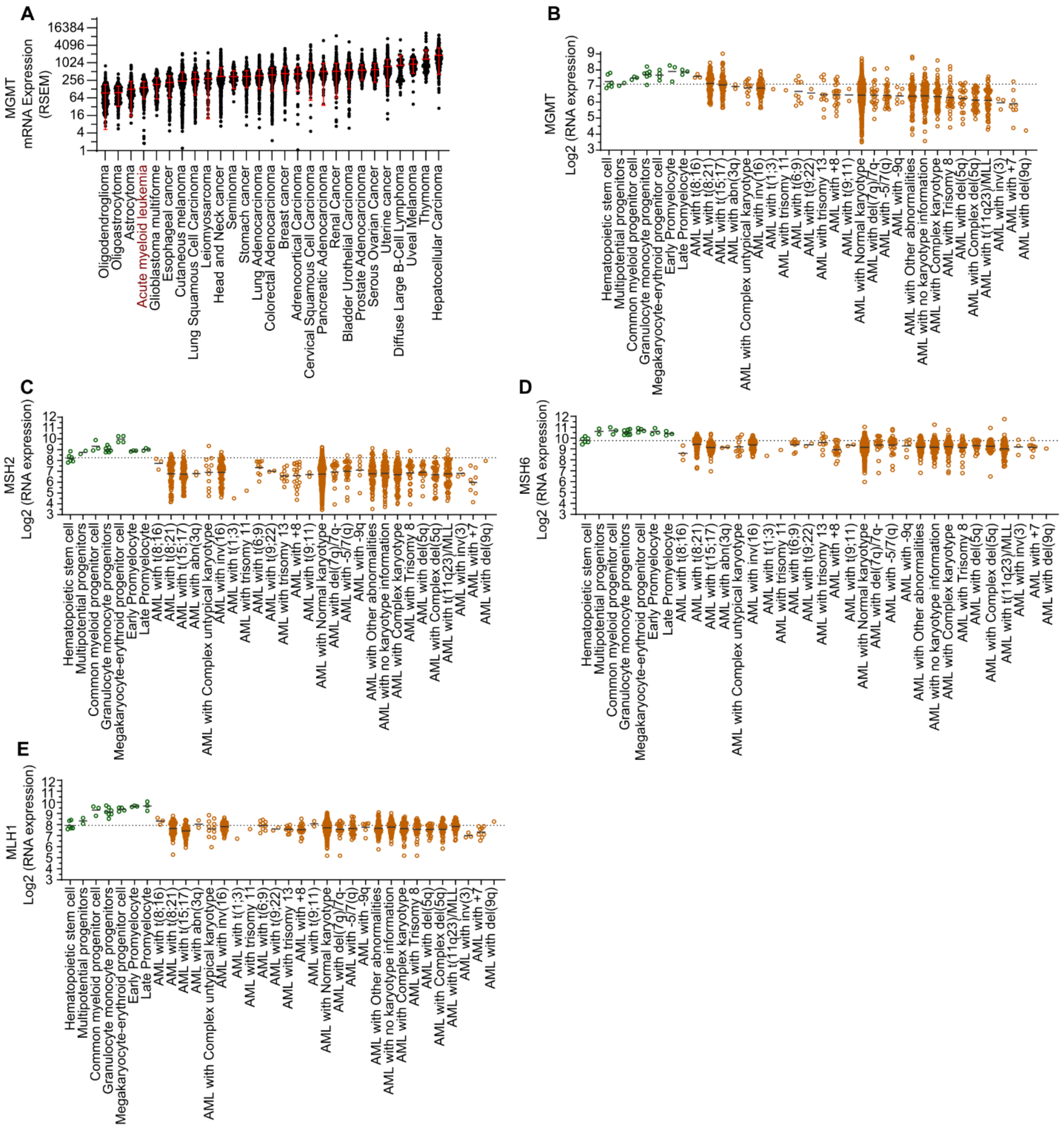


**Supplementary Figure 1: A)** MGMT mRNA expression levels across a pan-cancer cohort (Y-axis shown on a $Log_2$ scale. **B-E)** Batch-normalized **B)** MGMT, **C)** MSH2, **D)** MSH6, and **E)** MLH1 mRNA expression analysis comparing healthy donor hematopoietic stem and progenitor cells with various AML subtypes. Data were adapted from the BloodSpot database (ref 91), which provides integrated, batch-corrected RNA-seq results originally sourced from GSE42519, GSE13159, GSE15434, GSE61804, GSE14468, and the TCGA.

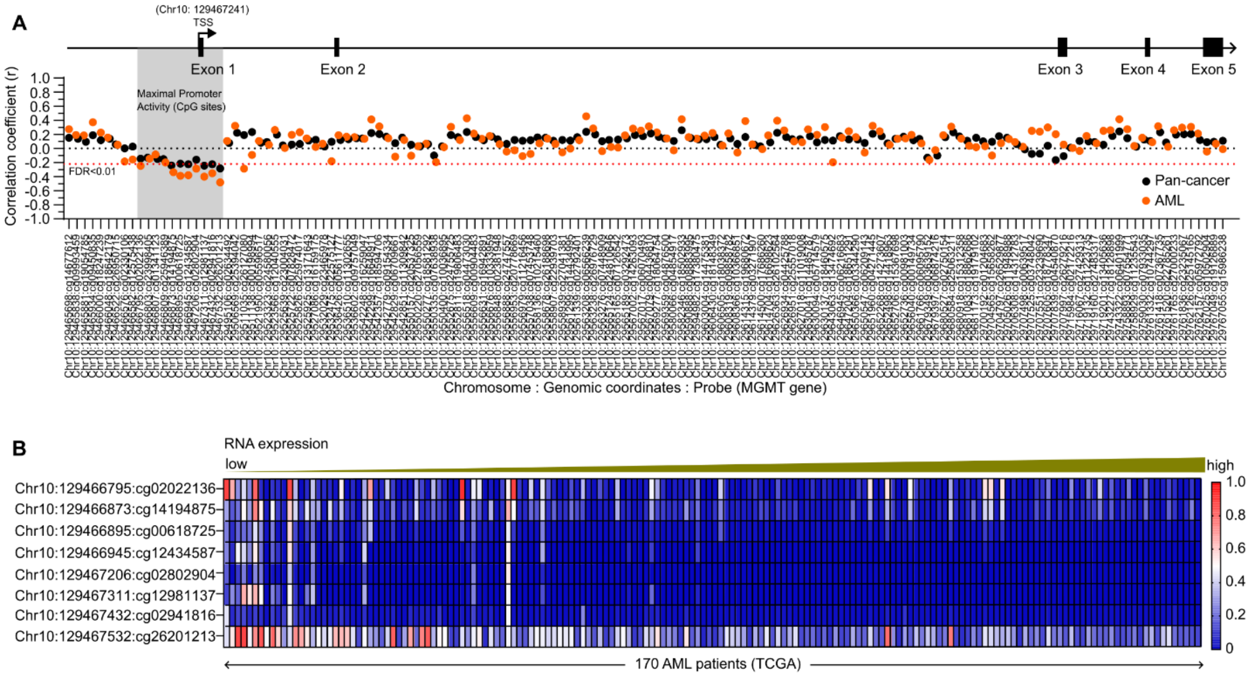


**Supplementary Figure 2: A)** Pearson correlation analysis of DNA methylation (beta-values) at specific MGMT loci with mRNA expression in the TCGA AML cohort (n=170). Analysis included 146 Illumina Human Methylation 450K probes spanning the MGMT gene, comparing AML (orange) against pan-cancer datasets (black). TSS represents the Transcription start site and the orange horizontal line denotes the FDR-adjusted significance threshold (q < 0.01; Benjamini-Hochberg procedure). **B)** Heatmap depicting methylation levels across eight statistically significant (q < 0.01) Infinium Methylation EPIC v2.0 probes or promoter sites, with patients (n=170) ranked by increasing MGMT mRNA expression.

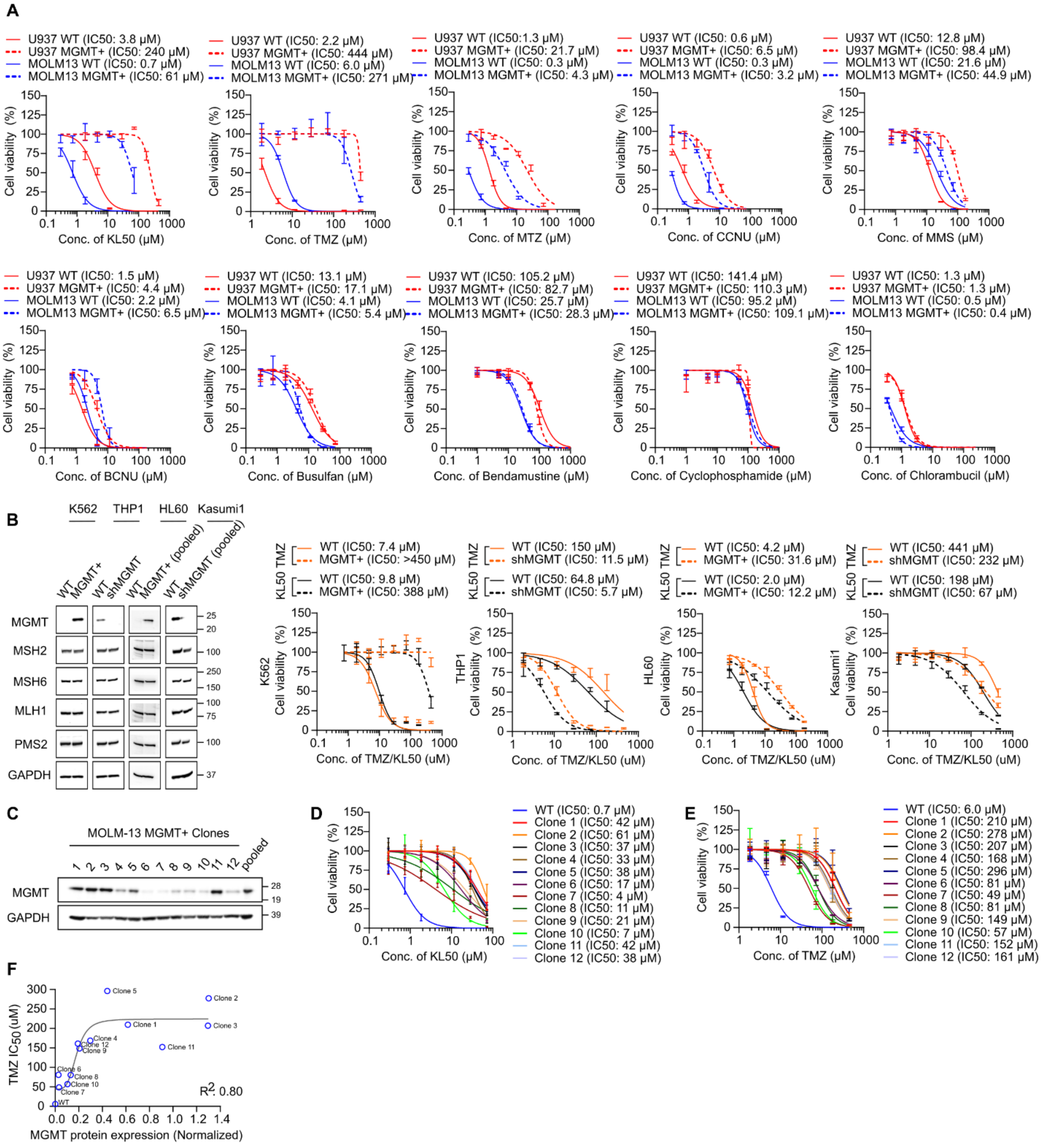


**Supplementary Figure 3: A)** Short-term *in vitro* viability assays evaluating diverse DNA alkylating agents in MGMT-isogenic pairs of MOLM13 and U937 cells. Abbreviations: TMZ, Temozolomide; KL50, 2-fluoroethyl-analog of TMZ; MTZ, Mitozolomide; CCNU, Lomustine; MMS, Methyl methanesulfonate; BCNU, Carmustine. **B)** Western blot confirming MGMT and MMR protein levels across engineered AML models following MGMT ORF-mediated over-expression (K562 and HL60) or shRNA-mediated MGMT knockdown (THP-1 and Kasumi-1). Corresponding dose-response curves illustrate the sensitivity of these models to KL50 and TMZ in short-term *in vitro* viability assays. **C-E)** Western blot analysis of MGMT expression across a panel of isogenic MOLM13 clones, with associated *in vitro* viability assays evaluating response to KL50 and TMZ. **F)** Correlation between MGMT protein levels and sensitivity to TMZ in isogenic MOLM13 clones. Goodness of fit was determined using a sigmoidal 4PL non-linear regression model.

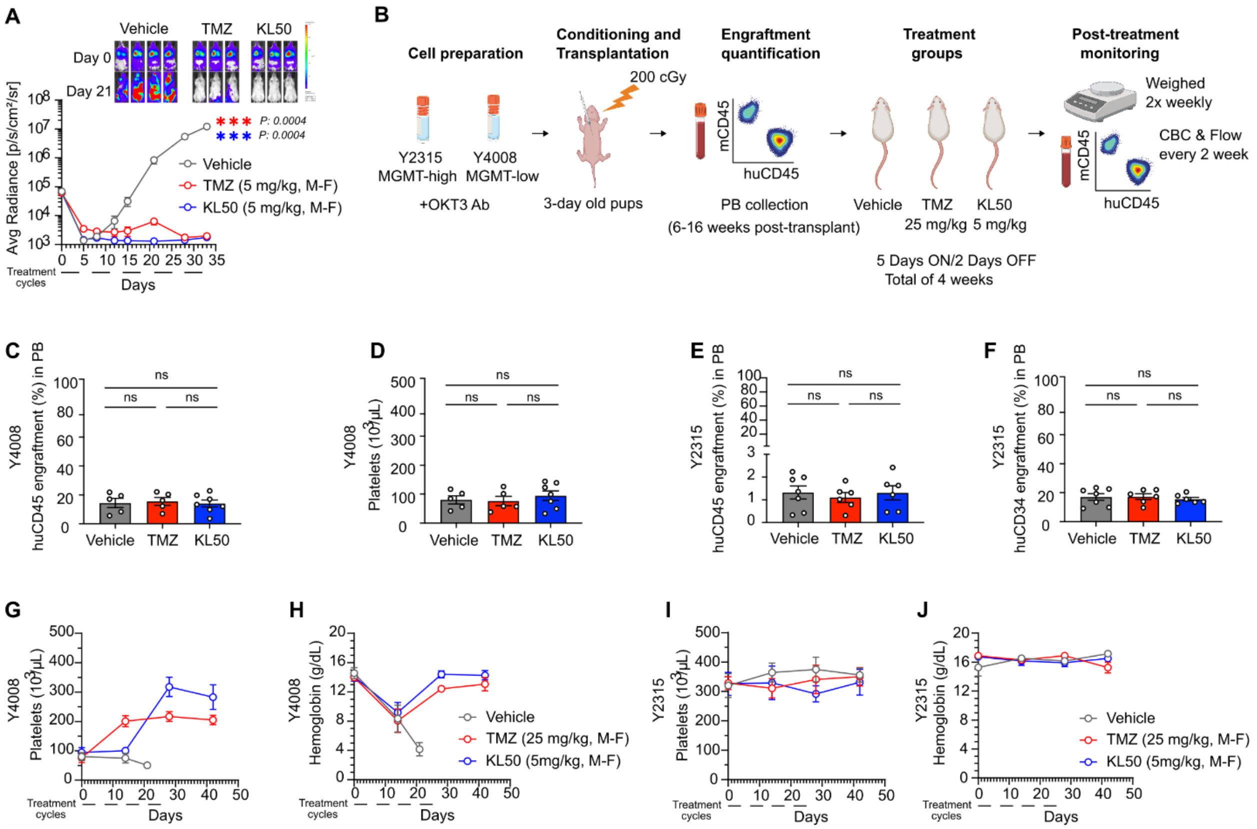


**Supplementary Figure 4: A)** *In vivo* evaluation of TMZ and KL50 in an intravenously (i.v.) engrafted MGMT⁻ U937 AML model using NSG (NOD-scid IL2Rgammanull) mice (Vehicle, n=4; TMZ, n=3; KL50, n=3). Inset: Representative bioluminescence imaging (BLI) of mice at Day 0 and Day 21 following i.v. engraftment of luciferase-expressing U937 cells. **B)** Schematic representation of primary AML sample engraftment and *in vivo* studies with TMZ and KL50 in MISTRG6 mice. **C-D)** Randomization of Y4008 secondary engrafted mice based on **C)** huCD45⁺ and **D)** platelets counts. **E-F)** Randomization of Y2315 secondary engrafted mice based on **E)** huCD45⁺ and **F)** huCD34⁺ levels. **G)** peripheral blood platelet counts, and **H)** hemoglobin levels of MGMT⁻ (Y4008) engrafted MISTRG6 mice following TMZ and KL50 treatment. **I)** peripheral blood platelet counts, and **J)** hemoglobin levels of MGMT⁺ (Y2315) engrafted MISTRG6 mice following TMZ and KL50 treatment. Treatment cycle: 5 days a week (M-F, Monday to Friday) for 4 weeks.

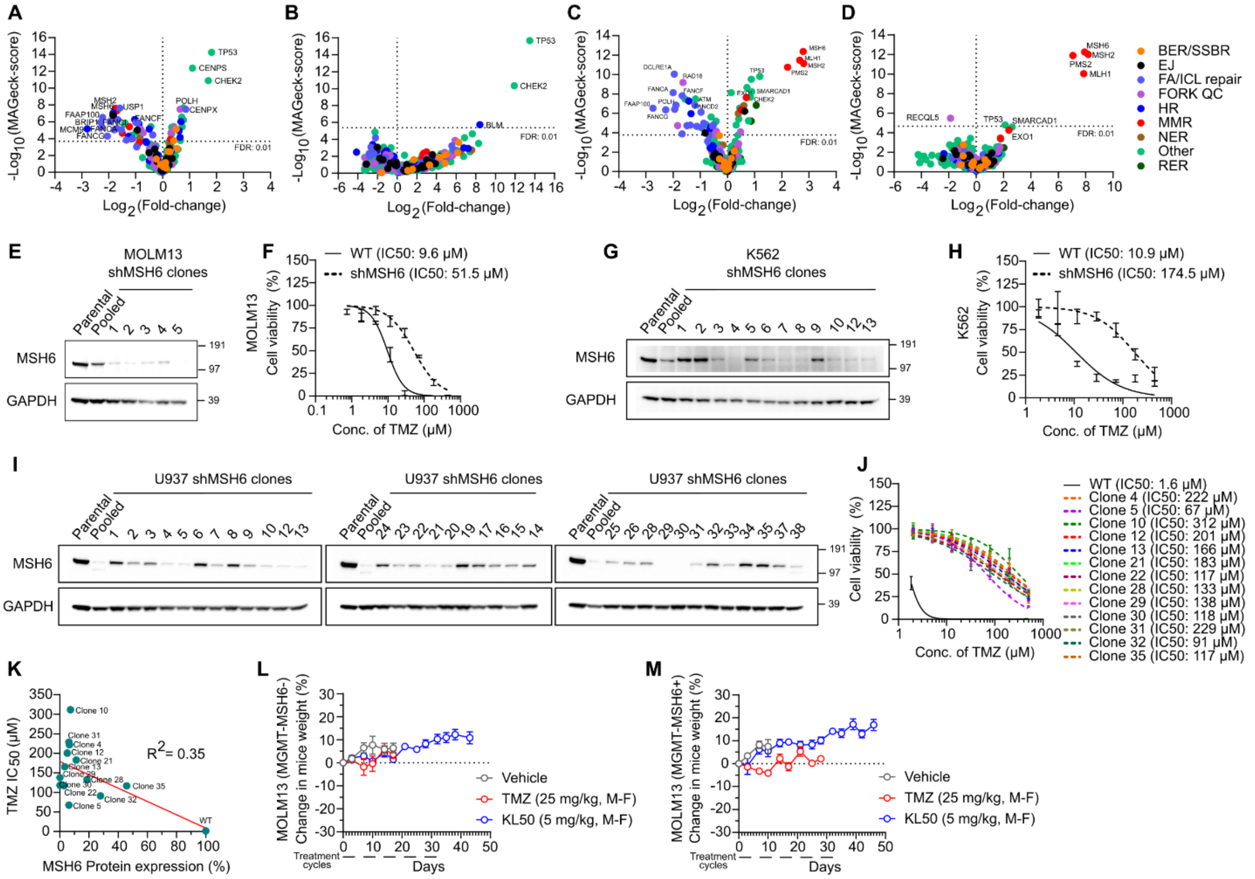


**Supplementary Figure 5: A-D)** DNA Damage Response (DDR)-focused CRISPR-knockout screen in parental MGMT- MOLM13 cells treated with **A)** 1.84 μM KL50, **B)** 7.36 μM KL50, **C)** 17 μM TMZ, and **D)** 46 μM TMZ. **E)** Western blot confirmation of reduced MSH6 expression following shRNA-mediated knockdown in MOLM13 clones. **F)** Short-term (6-day) *in vitro* viability assays with TMZ post shRNA knockdown of MSH6 in MOLM13 (shMSH6 clone 5). **G)** Western blot confirmation of reduced MSH6 expression following shRNA-mediated knockdown in K562 clones. **H)** Short-term (6-day) *in vitro* viability assays with TMZ post shRNA knockdown of MSH6 in K562 (shMSH6 clone 4). **I)** Western blot analysis of MSH6 protein levels and **J)** corresponding short-term *in vitro* viability assays with TMZ across a panel of shRNA-mediated MSH6-knockdown clones in U937 cells. **K)** Pearson correlation analysis between relative MMR deficiency and TMZ resistance across a panel of isogenic MSH6-knockdown U937 clones. MSH6 protein expression is normalized with GAPDH control and percentage was calculated in context with U937 WT cells. **L-M)** Longitudinal total body weight analysis of Athymic Nude (Foxn1nu) mice bearing subcutaneous **L)** MGMT- MSH6- and **M)** MGMT-MSH6+ MOLM13 xenografts during treatment with TMZ or KL50. Treatment cycle: 5 days a week (M-F, Monday to Friday) for 4 weeks.

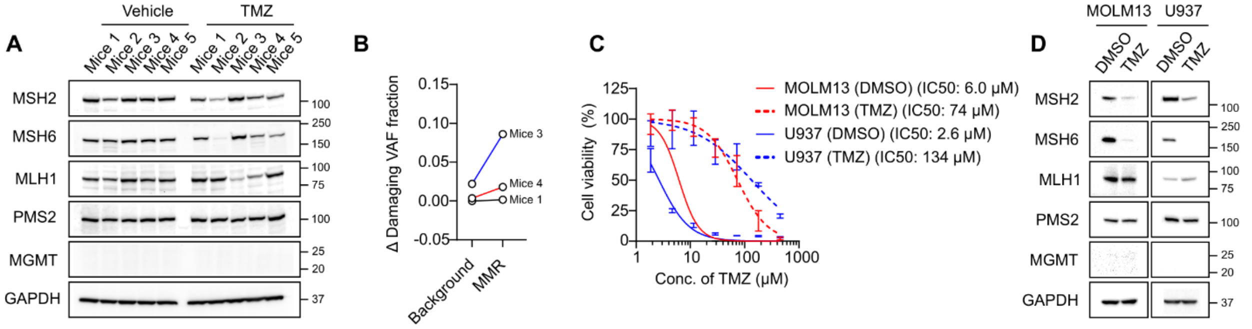


**E**

| Cells | Gene | Chromosome position | Damaging single nucleotide variant (SNV) | CADD score | Variant Allele Frequency (VAF) |
|---|---|---|---|---|---|
| MOLM13 | MSH2 | chr2:47471035 | G>T:missense_variant | 28.2 | 24% |
| U937 | MSH6 | chr2:47791119 | T>G:stop_gained | 35 | 33% |
| | | chr2:47801003 | G>A:stop_gained | 41 | 43% |

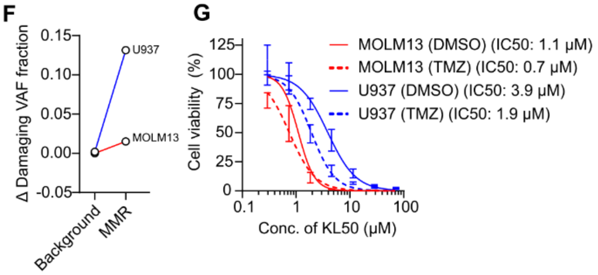


**Supplementary Figure 6: A)** Western blot analysis of harvested MOLM13 tumor cells evaluating MMR and MGMT expression post-relapse. **B)** WES analysis showing a change in damaging variant allele frequency (VAF) fraction in MMR genes as compared to the entire genomic background post 4-weeks TMZ treatment of representative MOLM13 tumors (n=3). VAF fractions were calculated by comparing TMZ-treated samples with their respective DMSO-treated controls. **C)** Short-term *in vitro* viability assays confirming the induction of TMZ resistance in AML cell lines following prolonged, escalating TMZ exposure. **D)** Western blot analysis revealing reduced MMR protein expression in MOLM13 and U937 cells post emergence of TMZ resistance. **E)** Table showing genomic coordinates, type of variant, allele frequency, and Combined Annotation Dependent Depletion (CADD) score of the damaging variant alleles detected in the MMR genes of MOLM13 and U937 cells post chronic TMZ treatment. **F)** WES analysis of TMZ-treated MOLM13 and U937 cells showing an increase in damaging VAF fraction in MMR genes over genomic background. **G)** Short-term *in vitro* viability assays demonstrating persistent KL50 sensitivity in cells with acquired MMR deficiency post TMZ.

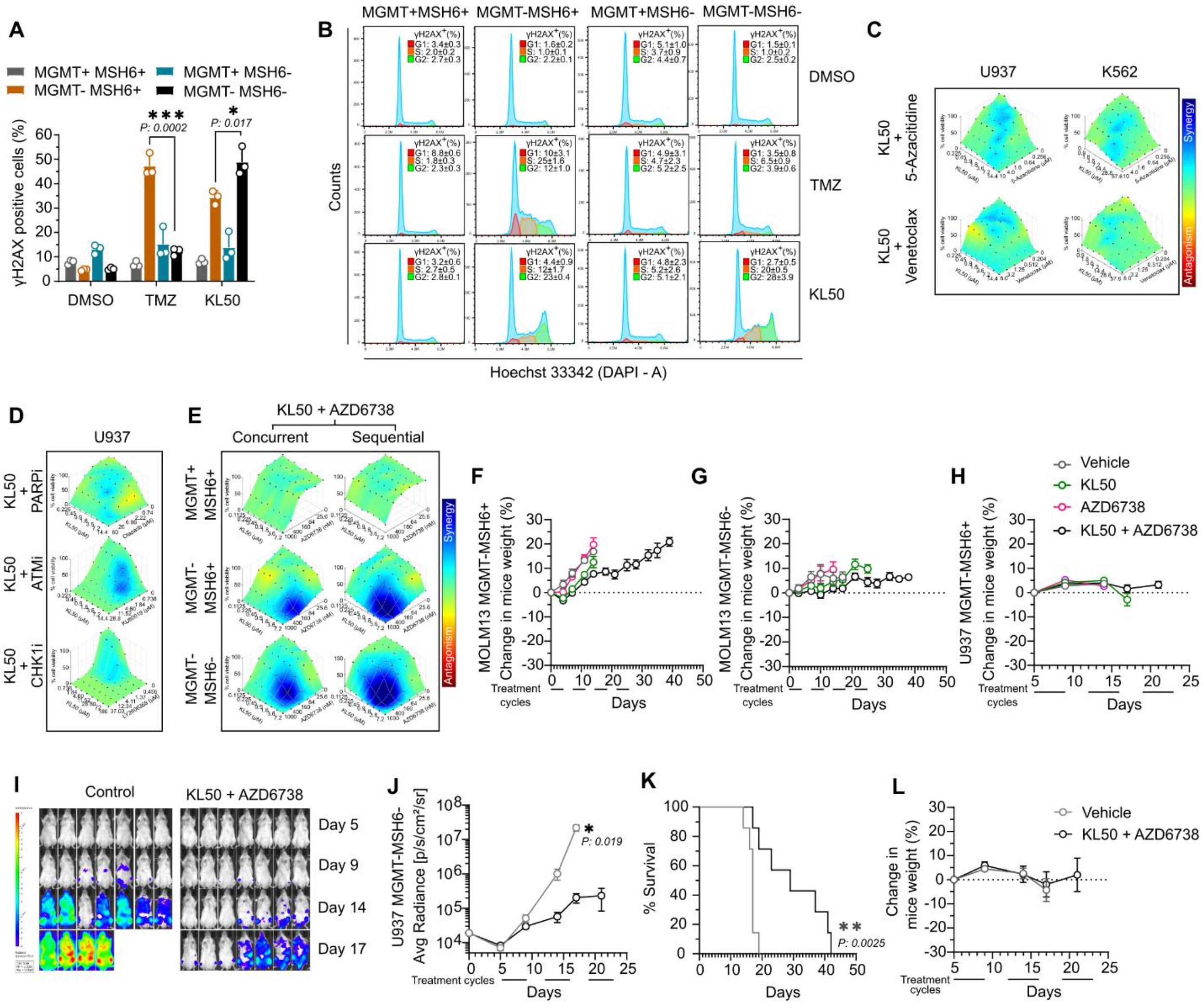


**Supplementary Figure 7: A)** Flow cytometry-based quantification of ɣH2AX levels in MGMT and MMR-isogenic U937 cells. **B)** Representative flow cytometry histograms depicting ɣH2AX induction across specific cell-cycle phases in the same MGMT and MMR-isogenic U937 cells. Inset: Quantification of the percentage of ɣH2AX-positive cells (mean±SD) across G1, S, and G2/M phases. **C)** *In vitro* synergy analysis (HSA synergy) between KL50 and AML standard of care regimens like 5-azacitidine or venetoclax in MGMT-silenced U937 and K562 parental cell lines. **D)** *In vitro* synergy between KL50 and PARP inhibitor (Olaparib), ATM inhibitor (KU60019), and CHK1 inhibitor (Prexasertib, LY2606368) in MGMT-silenced U937 parental cell line. **E)** Comparative *in vitro* synergy analysis of concurrent versus sequential dosing schedules of KL50 and ATRi in U937 isogenic cells. **F-H)** Longitudinal total body weight analysis of **F)** MGMT$^{-}$MSH6$^{+}$ MOLM13, **G)** MGMT$^{-}$MSH6$^{-}$ MOLM13 (subcutaneous), and **H)** MGMT$^{-}$MSH6$^{+}$ U937 (i.v. engrafted) xenograft-bearing mice during KL50/ATRi combination therapy. **I-L)** Characterization of the MGMT$^{-}$MSH6$^{-}$ i.v.engrafted U937 model following combination therapy, including **I)** representative bioluminescence imaging (BLI), **J)** quantitative flux analysis, **K)** Kaplan-Meier survival curves, and **L)** longitudinal body weight monitoring. Treatment cycle: Twice-weekly alternating doses (KL50: Mon/Thu; ATRi: Tue/Fri) for 3-4 weeks.
